# Sexually satiated male uses gustatory-to-neuropeptide integrative circuits to reduce time investment for mating

**DOI:** 10.1101/088724

**Authors:** Woo Jae Kim, Seung Gee Lee, Anne-Christine Auge, Lily Yeh Jan, Yuh Nung Jan

**Affiliations:** Department of Cellular and Molecular Medicine, University of Ottawa, Ottawa, Ontario K1H 8M5, Canada; Howard Hughes Medical Institute, Departments of Physiology, Biochemistry and Biophysics, University of California San Francisco, San Francisco, California 94158, USA

## Abstract

Males rely on a ‘time investment strategy’ to maximize reproductive success. Here we report a novel behavioral plasticity whereby male fruit flies exhibit a shortened mating duration when sexually satiated, which we named ‘Shorter-Mating-Duration (SMD)’. SMD requires the sexually dimorphic Gr5a-positive neurons for detecting female body pheromones. The memory circuitry within the ellipsoid body (EB) and mushroom body (MB) brain regions is crucial for SMD, which depends on the circadian clock genes *Clock* and *cycle*, but not *timeless* or *period*. SMD also relies on signaling via the neuropeptide sNPF, but not PDF or NPF. Sexual experience modifies the neuronal activity of a subset of sNPF-positive neurons involved in neuropeptide signaling, which modulates SMD. Thus, our study delineates the molecular and cellular basis for SMD – a plastic social behavior that serves as a model system to study how the brain switches the internal states between sexual drive and satiety.

## INTRODUCTION

From simple behaviors to sophisticated decisions, animals must make choices throughout their life in order to maximize their resources (Louâpre, van Alphen and Pierre, 2010). The reproductive success of a male animal is a function of the number of his sperm cells that are successful in fertilizing female eggs. Hence, sexual selection involves not just adaptive phenotypes for mating with many female partners (Parker, 1984). Males have a finite resource to spend on reproduction (Parker and Pizzari, 2010), which is limited by the number of ejaculates that can be delivered and the time required to restore depleted reserves (Dewsbury, 1982). Sperm can be costly to produce therefore males strategically allocate their investment into the ejaculate (Dowling and Simmons, 2012).

Besides ejaculate economics, time waste also can constitute a considerable selective disadvantage for males due to increased exposure to environmental hazards such as predators, or a lower fertilization rate, than their competitors. Regarding adaptation, time waste can be defined as where an individual spends longer than other animals to complete a given activity. Those individuals which waste time might expose themselves to the action of predators or various environmental hazards, then eventually fall into less competitive situations. In a species with a continuous life history. Moreover, time waste in males can result in direct disadvantages of the species by increasing the generation interval (Parker, 1974).

In a competitive mating environment, time-wasting males will have a significantly lower fertilization rate than their rivals. For males, the balance between the time investment for feeding and searching for females is the critical factor to increase their reproductive fitness (Parker, 1970). In this regard, a ‘time investment strategy (optimum allocation of time spent on given activities to achieve the maximum reproductive success)’ is crucial for males.

In support of the male’s time investment strategy, recent studies have revealed that *D. melanogaster* males vary greatly in their level of interest in females, providing evidence that males have also evolved mate selectivity behavior (Gowaty, Steinichen and Anderson, 2003). When mating opportunities are constrained, males show a preference for more fecund females, and in turn will benefit directly by increasing the number of offspring they produce (Byrne and Rice, 2006a). The stringent mating investment by *Drosophila* males might have evolved for the following reasons. First, sexual activity reduces the lifespan of males (Partridge and Farquhar, 1981) due to costs arising from vigorous courtship (Cordts and Partridge, 1996), the production of ejaculates (Lefevre and Jonsson, 1962), and possibly immunosuppression (McKean and Nunney, 2001). Second, repeated mating by males within a day depletes limiting components of the ejaculate (Demerec and Kaufman, 1941; Lefevre and Jonsson, 1962). Third, the quality of potential female mates is highly variable (Lefranc and Bundgaard, 2000). Thus, male flies’ efforts for mating might vary in different contexts.

Behavioral plasticity is preferable when specific aspects of the environment (e.g., the intensity of socio–sexual encounters) are prone to rapid and unpredictable variation (Bretman, Gage and Chapman, 2011). Behavioral plasticity requires the formation of association between a proper behavioral output and given information, as the consequences of multiple interactions between evolutionarily programmed innate behaviors and cumulated learning experiences of the animal (Dissel, Angadi, Kirszenblat, Suzuki, Donlea, Klose, Koch, English, Winsky-Sommerer, van Swinderen and Shaw, 2015; Lupold, Manier, Ala-Honkola, Belote and Pitnick, 2011).

What are the general terms of behavioral plasticity and what is its meaning in the context of the fruit fly’s sexual behavior? One example of plastic male behavior is ‘Longer-Mating-Duration (LMD)’, with increased investment via mating duration lengthening induced by exposure to rivals before mating (Bretman, Fricke and Chapman, 2009). In *Drosophila,* males respond to the presence of rivals by prolonging mating duration to guard the female and pass their genes. In previous studies, we examined the genetic network and neural circuits that regulate rival-induced longer mating duration (LMD). LMD can be induced solely via visual stimuli. LMD depends on the circadian clock genes *timeless* and *period*, but not *Clock* or *cycle*. LMD involves the memory circuit of the ellipsoid body (EB). Further, we identified a small subset of clock neurons in the male brain that regulates LMD via neuropeptide signaling (Kim, Jan and Jan, 2013).

Here we report a novel plastic behavior of male *D. melanogaster* for its selective investment in mating. Sexually satiated *Drosophila* males show this plastic behavior by limiting their investment in copulation time, namely ‘Shorter-Mating-Duration (SMD)’. In addition to delineating the requisite circuitry by identifying the neurons for SMD, our study has uncovered the sensory stimuli, clock genes, sexual dimorphism, and neuropeptide signaling crucial for the SMD behavior, which is functionally and mechanistically distinct from the previously identified LMD pathway.

## RESULTS

### Sexually satiated males reduce investment in mating duration as compared to naïve males

To investigate how sexual experience affects the mating duration of male fruit flies, we introduced virgin females into group-reared males 1 day before the assay (this condition will be referred as ‘experienced’ henceforward), and compared their mating duration with group-reared males that never encountered sexual experience (this condition will be referred as ‘naïve’ henceforward) (Figure 1A). We found that the mating duration of wild-type naïve males such as Canton-S (*CS*), WT Berlin (*WT^Ber^*), and Oregon R (*Ore^R^*) is significantly longer than that of sexually experienced males (Figure 1B). To test whether the genetic background of male flies affect the SMD behavior, we performed mating duration assay with laboratory maintained *w^1118^* males. We found in our previous study that *w^1118^* males do not show the LMD behavior as a plastic responses of male fruit flies to the level of sperm competition, which results in significantly increased reproductive success in a competitive environment by extending mating duration (Bretman, Fricke and Chapman, 2009), probably because *w^1118^* males have defective vision (Kim et al., 2013). In contrast, we found the SMD behavior was normal in *w^1118^* mutant (*w^1118^* in Figure 1B). We also found that socially isolated males show SMD behavior (Figure S1A). Thus, unlike LMD, SMD persists in either grouped or isolated rearing conditions. Therefore, we decided to perform all mating duration assays with a group-reared condition.

**Figure 1.**
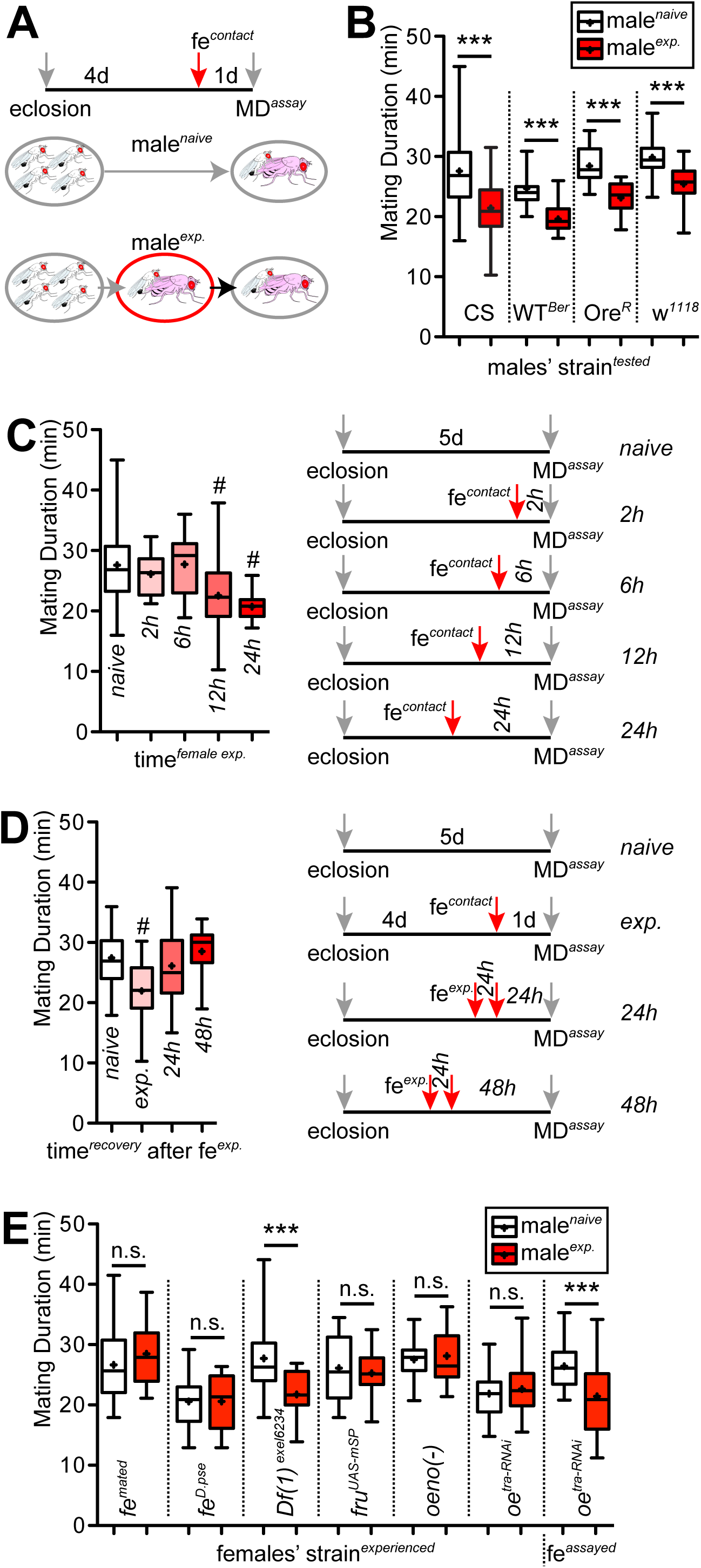
General characteristics of ‘Shorter-Mating-Duration (SMD)’ behaviour. (A) Naïve males were kept for 5 days after eclosion in groups of 4 males. Experienced males were kept for 4 days after eclosion in groups then experienced with 5 virgin females 1 day before assay; for detailed methods, see **EXPERIMENTAL PROCEDURES**. (B) Mating duration (MD) assays of Canton-S (CS), Oregon-R, WT-Berlin and *w^1118^* males. White boxes represent naïve males and grey boxes represent experienced ones. (C) MD assays of CS males experienced with females in different time duration. Different female-experienced times are described below the boxes. (D) MD assays of CS males after isolated from female experience. Males were reared with sufficient numbers of virgin females for 2 h to be assured having sexual experience then isolated. Assay times after isolation are described below the boxes. Boxes labeled naïve and exp. represent standard rearing conditions described in **Figure 1A.** (E) MD assay of CS males experienced with different female strains 1 day before assay. Experienced females’ genotypes are described below the boxes. To make mated females (*fe^mated^*), 4-day-old 10 CS virgin females were placed with 5-day-old 20 CS males for 6 hours then separated to an empty vials. These females were used for experienced females 1 day after separation. *fe^D.Pse^* is *D. pseudoobscura* females. *Df^exel6234^* is deficiency strain that lacks *sex-peptide receptor* (SPR) expression, the critical component for post-mating switch in female *Drosophila (Yapici, Kim, Ribeiro and Dickson, 2008)*. To make virgin females behave as mated females (*fru^UAS-mSP^*), flies expressing *UAS-mSP* (membrane bound form of male sex-peptide) were crossed with flies expressing *fru-GAL4* driver as described previously (Yang, Rumpf, Xiang, Gordon, Song, Jan and Jan, 2009). To make oenocyte-deleted females (*oenocyte (-)*), flies expressing *UAS-Hid/rpr* virgins were crossed with flies expressing *tub-GAL80ts; oeno-GAL4* males then the female progenies were kept in 22° for 3 days. Flies were moved to 29° for 2 days before assay to express *UAS-Hid/rpr* to kill the oenocytes in these females. The *oeno-GAL4* (*PromE(800)-GAL4*) was described previously (Wang, Han, Mehren, Hiroi, Billeter, Miyamoto, Amrein, Levine and Anderson, 2011b). To make oenocytes-masculinized females (*oeno^tra-RNAi^*), flies expressing *UAS-tra-RNAi* were crossed with *oeno-GAL4* driver. To test whether genotypes of female partners during MD assay affect MD, oenocyte masculinized females (*oeno^tra-RNAi^*) were used. Box plots represent the median value (horizontal line inside box), the mean value (‘+’ symbol inside box), interquartile range (height of the box, 50% of the data within this range), and minimum and maximum value (whiskers). Asterisks represent significant differences revealed by Student’s *t* test (* *p*<0.05, ** *p*<0.01, *** *p*<0.001). The same notations for statistical significance are used in other figures. Number signs represent significant differences revealed by Dunn’s Multiple Comparison Test (^#^ *p*<0.05). The same symbols for statistical significance are used in all other figures. See **EXPERIMENTAL PROCEDURES** for detailed statistical analysis used in this study.

We wondered whether the reduced mating duration of sexually experienced males may have been caused by the fatigue of repetitive sex. To test whether fatigue causes SMD behavior, we examined other behavioral repertories of naïve and experienced male flies, such as courtship index (a measure of the time a male engages in any defined recognizable courtship behaviors), courtship latency (time to courtship initiation), copulation latency (time to copulation/mating initiation), and locomotion (any of a variety of movements or methods that animals use to move from one place to another), and found no significant difference between experienced and naïve males (Figure S1B-G). Since several behavioral repertories of experienced males that should be affected by their fatigue were comparable to those of naïve males, we conclude that fatigue of repetitive female experiences is not a factor in causing SMD behavior.

Recently Zhang and colleagues reported that male flies show reproductive satiety when males are exposed to an excessive number of females. In their experimental paradigm, the percentage of time males spend in mating behaviors gradually declines to ~10% over 4 h (Zhang, Rogulja and Crickmore, 2016). This finding suggests that males do not invest much time for mating behaviors when they are satiated with females after 4 h of mating experiences. To test whether the SMD behavior is different from reproductive satiety, we designed a series of experiments by varying the time of male exposure to females. We found that males shortened their mating duration when their exposure to females lasted for 12 h rather than 6 h or 2 h, suggesting that SMD requires chronic exposure to females for longer than 6 h (Figure 1C). This suggests that, whereas 4 h of mating experience can induce reproductive satiety and reduce the mating drive, it is not sufficient to induce SMD behavior.

One of the crucial features of behavioral plasticity is the reversibility, which allows animals to adapt to fast changing environment. To determine whether SMD is a reversible behavior, we isolated males from females after 24 h of sexual experience then performed mating duration assay. We found that separating experienced males from females for 24 h was sufficient to restore the MD to the level of naïve males (Figure 1D). All these data suggest that SMD is plastic; it can change over time and may vary with the context, and is dependent on sexual experience with females.

Males should be especially prudent in the allocation of their limited resources such as the amount of sperms, which determines the reproductive success of male fruit fly (Lupold, Manier, Ala-Honkola, Belote and Pitnick, 2011). Given that sperm depleted males prefer large females as partners in courtship and copulation (Byrne and Rice, 2006b), we hypothesized that sperm depletion of experienced males may affect SMD behavior. To test the effect of sperm depletion on mating duration, we designed a series of experiments by varying the number of virgin females presented to a male, by up to a factor of 10. Because 5 h exposure with 4 females (Byrne and Rice, 2006b) or four consecutive copulation with females (Demerec and Kaufmann, 1941; Lefevre and Jonsson, 1962) depletes the majority of sperms, we decided to introduce 10 females to a male for a certain amount of time in order to deplete most of his sperm. We found that the mating duration of a male exposed to 10 females for 2 h, 4 h, or 8 h is comparable to that of control males (Figure S1H). To further confirm the lack of effect of sperm depletion on SMD behavior, we tested the son-of-*tudor* males that lack germ cells and therefore are spermless (Xue and Noll, 2000). Spermless son-of-*tudor* males show the intact SMD behavior suggesting that the absence of germ cells do not affect SMD behavior (Figure S1I). Consistent with our findings, Crickmore and colleagues have shown that the mating time of male fruit fly is not determined by the volume or rate of transfer of reproductive fluids (Crickmore and Vosshall, 2013). All these data suggest that sperm depletion of male does not cause SMD behavior.

To look for the physiological cues from females that produce SMD behavior, we first introduced various types of females as sexual partners to condition males so they would become “experienced”, before mating duration assay. We found that mated females and *D. pseudoobscura* virgin females cannot induce SMD behavior (*fe^mated^* and *fe^D.pse^* in Figure 1E). Mated females are known to display the post-mating responses (PMR) via sex peptide (SP)/sex peptide receptor (SPR) signaling and reject males (Yapici, Kim, Ribeiro and Dickson, 2008). It is also known, and confirmed in our experiments, that male *D. melanogaster* cannot mate with *D. pseudoobscura* females (data not shown). Next, we asked whether mating experiences with *D. melanogaster* females are crucial to producing SMD. We first tested SPR deficiency mutant females, *Df(1)^exel6234^*, that lack the SPR protein and remain receptive to males, exhibiting virgin-like behaviors after mating (Hasemeyer, Yapici, Heberlein and Dickson, 2009; Rezaval, Pavlou, Dornan, Chan, Kravitz and Goodwin, 2012; Yang, Rumpf, Xiang, Gordon, Song, Jan and Jan, 2009; Yapici, Kim, Ribeiro and Dickson, 2008). Males experienced with the sex-peptide receptor (SPR) mutant females exhibited the normal extent of SMD (*Df(1)^exel6234^* in Figure 1E). We then asked how SMD behavior may be affected if males were conditioned via exposure to virgin females behaving like mated ones. We found that SMD could not be induced by virgin females with expression of the membrane-bound form of male sex-peptide in *fruitless*-positive neurons, which behave like mated females that are not receptive to mating (Hasemeyer, Yapici, Heberlein and Dickson, 2009; Yang, Rumpf, Xiang, Gordon, Song, Jan and Jan, 2009; Yapici, Kim, Ribeiro and Dickson, 2008) (*fru^UAS-mSP^* in Figure 1E). These results suggest that the mating experiences are necessary to induce SMD behavior.

We next asked what kinds of physiological cues are provided by females to induce SMD behavior of males. Oenocytes are secretory cells that are the major sites synthesizing cuticular hydrocarbons including pheromones, which is critical to insect communication (Makki, Cinnamon and Gould, 2014). To test whether the cues produced from the oenocytes are essential to induce SMD, we produced pheromone-free females by ablating female oenocytes *(oeno(-)*). To test whether the female form but not the male form of cuticular hydrocarbons is required to generate SMD behavior, we generated females expressing a male-odor by masculinization of female oenocytes (*oe^tra-RNAi^*). We confirmed that these females show normal mating behavior with wild type males (data not shown). Males experienced with these females did not show SMD behavior (*oeno(-)* and *oe^tra-RNAi^* in Figure 1E), suggesting that female-specific pheromones produced in oenocytes are important cues for inducing SMD. If an oenocyte-masculinized female served as the mating duration assay partner rather than females providing mating experiences to males in the experienced group, the SMD remained normal (*fe^assayed^* in Figure 1E). This data suggests that male’s decision on mating duration heavily depends on the previous mating experiences, not the current one. All these findings indicate that the mating experience and the *D. melanogaster* female-specific odors produced by oenocytes are both required to induce SMD behavior.

### Contact chemoreception is critical for inducing SMD behavior

Male flies can detect females with various sensory modalities. In the early steps of courtship rituals, mainly visual, vibratory and olfactory signals allow males to find females and to orient towards females. In the next steps, males contact females by tapping with their forelegs to detect non-volatile hydrocarbon pheromones on the female cuticle via the gustatory system (Fernandez and Kravitz, 2013). To identify the sensory modalities that modulate SMD behavior, we tested the following mutants with defects on each sensory modality as described before (Kim, Jan and Jan, 2012, 2013). To test the visual contribution to SMD behavior without genetic intervention, we first performed an MD assay with animals reared in constant darkness for 5 days. SMD behavior was intact under constant dark conditions (*dark* in Figure 2A). Blind males with photoreceptors removed via *GMR-hid* also showed SMD (*GMR^Hid^* in Figure 2A). SMD was also normal in mutants with impaired vision, such as *ninaE^17^* without rhodopsins in R1-6 photoreceptors *(ninaE^17^* in Figure 2A) (Cook, Pichaud, Sonneville, Papatsenko and Desplan, 2003). Indeed, visually defective *w^1118^* mutant also displayed normal SMD behavior (Figure 1B). All these data suggest that vision is not important for SMD behavior.

**Figure 2.**
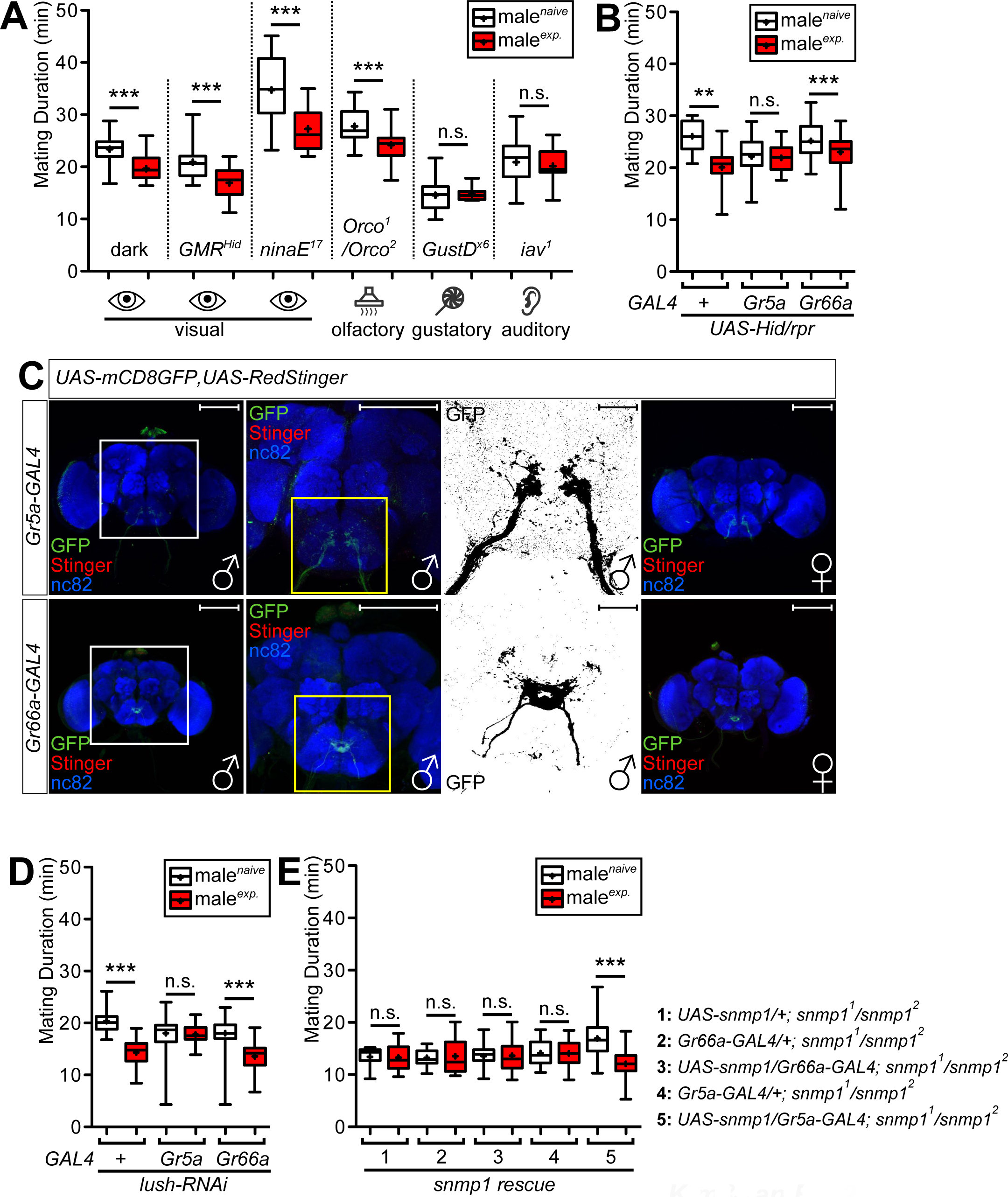
Pheromone sensing proteins expressed in *Gr5a*-positive neurons are responsible to induce SMD. (A) MD assay of various vision, olfactory, and auditory mutants. Genotypes are indicated below the graphs. To test vision is required for SMD, CS males were reared and sexually experienced in constant dark for entire 5 days (dark). (B) MD assays for *GAL4* driven cell death of which labels sweet cells (*Gr5a*) or bitter cells (*Gr66a*) using *UAS-Hid/rpr*. (C) Brains of flies expressing *Gr5a-GAL4 or Gr66a-GAL4* together with *UAS-mCD8GFP, UAS-RedStinger* were immunostained with anti-GFP (green), anti-DsRed (red) and nc82 (blue) antibodies. Scale bars represent 100 μm in the 1^st^, 2^nd^, and 4^th^ panels, 10 μm in the 3^rd^ panels from the left. White and yellow boxes indicate the magnified regions of interest presented next right panels. The 3^rd^ panels from the left are presented as grey scale for clearly showing the axon projection patterns of gustatory neurons in the adult subesophageal ganglion (SOG) labeled by *GAL4* drivers. (D) MD assays for *GAL4* mediated knockdown of LUSH via *UAS-lush-IR; UAS-dicer (lush-RNAi)*. Names of the *GAL4* drivers are indicated below the graphs. (E) Mating duration assays of *snmp1^1^* rescue experiments. Genotypes are indicated above the graphs.

To test the possible involvement of the olfactory pathway, we performed mating duration assay with olfactory mutants (*Orco^1^*/*Orco^2^)* with disrupted behavioral and electrophysiological responses to many odorants (Louis, Huber, Benton, Sakmar and Vosshall, 2008). Interestingly, the *Orco^1^*/*Orco^2^* trans-heterozygote mutant displayed normal SMD behavior (*Orco^1^*/*Orco^2^* in Figure 2A), suggesting that the olfactory pathway is not crucial for SMD induction. We then asked whether the gustatory pathway is important for SMD behavior. Gustatory mutants (*GustD^x6^)* with aberrant responses to sugar and NaCl (Rodrigues, Sathe, Pinto, Balakrishnan and Siddiqi, 1991) did not exhibit SMD behavior (*GustD^x6^* in Figure 2A), suggesting that the gustatory pathway is a critical sensory modality to induce SMD behavior. We also tested the auditory mutant *iav^1^* (Bretman, Westmancoat, Gage and Chapman, 2011) and found that they also did not exhibit SMD (*iav^1^* in Figure 2A). In summary, we found evidence for the involvement of gustatory and auditory pathways in generating SMD, although we cannot rule out the possible involvement of other sensory stimuli.

Flies touch one another upon their encounter. Taste and touch signals are conveyed to the brain by sensory neurons in the legs and mouthparts (Billeter and Levine, 2013). Male flies can detect pheromones by pheromone-sensitive cells that are incorporated into the chemosensory systems of taste and smell. We decided to focus on the taste system since we found that olfactory stimuli are not critical to induce SMD behavior (Figure 2). So we next asked whether we can identify the particular types of male taste neurons that are essential to detect female cues in eliciting SMD behavior. As reported previously (Wang, Singhvi, Kong and Scott, 2004), Gr5a- and Gr66a-positive neurons in the fly gustatory system correspond to two non-overlapping neuronal populations. Gr5a-positive cells mediate sweet-taste detection, whereas Gr66a-positive cells mediate bitter-taste recognition. We found that male flies with ablated Gr5a-positive neurons cannot elicit SMD behavior while male flies lacking Gr66a-positive neurons show normal SMD (Figure 2B), suggesting that Gr5a-positive neuronal populations are required for SMD. SMD was also impaired by by expressing Kir2.1 potassium channel in Gr5a-positive neurons in an adult-specific manner using *tub-GAL80^ts^* (Figure S2A). To identify those neurons that express Gr5a or Gr66a, we characterized the expression pattern driven by the *Gr5a-GAL4* or *Gr6a-Gal4* driver using the reporter *UAS-mCD8-GFP* (a cell membrane marker) along with *UAS-RedStinger* (a nuclear marker). We confirmed that these two neuronal populations have non-overlapping axonal projections to the suboesophageal ganglion (SOG) region in both males and females (Figure 2C) (Wang, Singhvi, Kong and Scott, 2004). All these results confirm that the Gr5a taste neurons are essential for generating SMD behavior.

Pheromones and the mechanisms for their detection have evolved to transmit biologically relevant information from one member of a species to another (Mucignat-Caretta, 2014). Whereas the olfactory system is important for volatile pheromone communications, insect gustatory neurons present on the tarsi (legs) are also required for courtship behavior, and are selective for the detection of male or female cuticle lipids, to mediate ‘contact chemoreception’ (Mucignat-Caretta, 2014). LUSH is an odorant-binding protein (OBP) expressed in the antenna, and is also found in a subset of tarsal chemoreceptors on the forelegs (Kim, Repp and Smith, 1998). We asked whether the expression of LUSH in Gr5a-positive neurons is responsible for pheromone signaling of SMD induction. Knocking down the expression of LUSH in Gr5a-positive neurons diminished the SMD behavior, whereas knocking down LUSH expression in Gr66a-positive neurons had no effect on SMD (Figure 2D), suggesting that LUSH expression in Gr5a-positive neurons are crucial for detecting female pheromones for producing SMD behavior.

Snmp1 is in the CD36 related protein family and functions as an important player for the rapid kinetics of the pheromone response in insects (Li, Ni, Huang and Montell, 2014). We found that various genotypes of *snmp1* mutants did not show SMD behavior (Figure S2B), suggesting that expression of Snmp1 protein is necessary for SMD behavior. Snmp1 is prominently expressed in olfactory sensory neurons in the antenna, and is also expressed in chemosensory organs on the proboscis (Benton, Vannice and Vosshall, 2007). We found that expression of Snmp1 in the *snmp1* mutant background via the *Gr5a-GAL4* driver could rescue SMD behavior (Figure 2E), suggesting that expression of Snmp1 protein only in Gr5a-positive neurons is sufficient to induce SMD behavior. Taken together, these data suggest that contact chemoreception mediated by the pheromone sensing proteins LUSH and Snmp1 in Gr5a-positive gustatory neurons is critical for triggering SMD behavior.

### Male-specific Gr5a-positive neurons are required for SMD

Mating duration of *D. melanogaster* is solely determined by male and is associated with significant fitness benefits (Bretman, Westmancoat and Chapman, 2013). We previously showed that a small number of male-specific neurons control the rival-induced extended mating duration, LMD behavior (Kim, Jan and Jan, 2013). Since SMD behavior is also associated with modulation of mating duration, we hypothesized that male-specific and sexually dimorphic neural circuits modulate this behavior. It is well known that sexual dimorphism of sensory structure and function generates neuronal circuitries important for gender-specific behaviors (Zarkower, 2001). In *Drosophila*, *fruitless (fru)* is an essential neural sex-determinant responsible for male-specific behaviors (Ryner, Goodwin, Castrillon, Anand, Villella, Baker, Hall, Taylor and Wasserman, 1996). To determine whether sexually dimorphic sensory neurons are involved in SMD, we used intersectional methods to genetically dissect ~1500 *fru* neurons into smaller subsets. We used a combination of the *fru^FLP^* allele that drives FLP-mediated recombination specifically in *fru* neurons with *UAS[stop]X* (X could be various reporters or effector transgenes) to express a *UAS* transgene in those cells that are not only labeled by the *GAL4* driver but are also *fru*-positive, due to FLP-mediated excision of the stop cassette (*[stop]*) (Yu, Kanai, Demir, Jefferis and Dickson, 2010a).

We first asked if there are *fru*-positive and sexually dimorphic sensory neurons among *Gr5a*-positive cells. When we used *Gr5a-GAL4* to identify sexually dimorphic Gr5a-positive cells, we found a restricted numbers of FRU-positive and Gr5a-positive neurons in the male forelegs and proboscis (compare GFP to Stinger in top panels of Figure 3A). In contrast, we could not find any FRU-positive and Gr5a-positive neurons in female foreleg and proboscis (middle panels in Figure 3A), suggesting that these FRU-positive and Gr5a-positive sensory neurons are sexually dimorphic. We failed to label FRU-positive neurons in male forelegs and proboscis using *Gr66a-GAL4* driver (bottom panels in Figure 3A), suggesting that there are no FRU-positive and Gr66a-positive sensory neurons in male’s forelegs and proboscis. Next we asked whether these male-specific neurons project their axons into the SOG (suboesophageal ganglion) region receiving gustatory inputs. We confirmed the axonal projections of these male-specific Gr5a-positive neurons to the SOG region with the membrane marker *UAS-mCD8GFP* (Figure 3B) and the presynaptic marker *UAS-nsybGFP* (Figure 3C). We could not find any GFP signals with this manipulation within the female brain (Figure S3A). The same genetic manipulation to label FRU-positive and Gr5a-positive neurons with the dendritic marker *UAS-DscamGFP* failed to label Gr5a-positive GFP signals within the SOG region (Figure S3B), suggesting that the GFP fluorescence shown in Figure 3B corresponds to presynaptic axon terminals, instead of dendrites. All these data suggest that sexually dimorphic Gr5a-positive and FRU-positive neurons in male forelegs and proboscis project their axons into the SOG region of the male brain.

**Figure 3.**
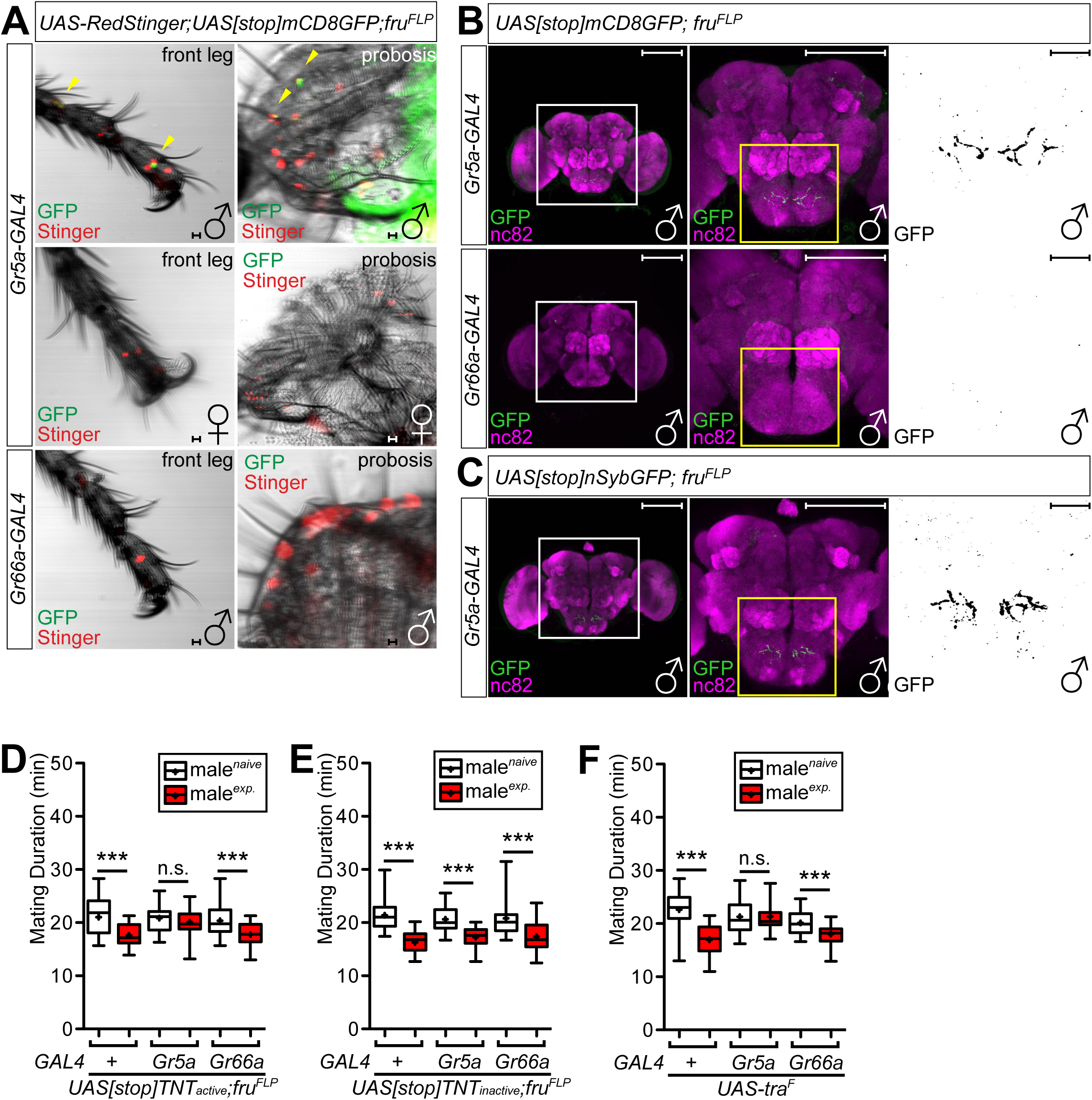
Male specific FRU-positive subsets of *Gr5a*-positive neurons are required to generate SMD. (A) Front legs and proboscis of flies expressing *Gr5a-* and *Gr66a-GAL4* drivers together with *UAS-RedStinger; UAS[stop]mCD8GFP; fru^FLP^* were imaged live under fluorescent microscope. Yellow arrows indicate *fruitless-*positive neurons. Scale bars represent 10 μm. (B) Brains of flies expressing *Gr5a-GAL4 or Gr66a-GAL4* together with *UAS[stop]mCD8GFP; fru^FLP^* were immunostained with anti-GFP (green) and nc82 (magenta) antibodies. White and yellow boxes indicate the magnified regions of interest presented next right panels. The right panels are shown grey scale for clearly presenting the axon projection patterns of gustatory neurons in the adult SOG labeled by *GAL4* drivers. Scale bars represent 100 μm. (C) Same experiment of **Figure 3B** using presynaptic marker *UAS[stop]n-sybGFP; fru^FLP^*. (D) MD assays of *Gr5a-* and *Gr66a-GAL4* drivers for inactivation of synaptic transmission of male-specific neurons among each *GAL4*-labeld neurons via *UAS[stop]TNTactive; fru^FLP^*. (E) Control experiments of **Figure 3D** with inactive form of *UAS-TNT using UAS[stop]TNTinactive; fru^FLP^*. (F) Mating duration assays for *Gr5a-* and *Gr66a-GAL4* drivers for feminization of neurons via *UAS-tra^F^*.

We next analyzed the function of these male-specific neurons on SMD behavior. We first confirmed that inhibiting synaptic transmission by TNT expression via *Gr5a-GAL4* but not *Gr66a-GAL4* specifically eliminated SMD behavior (Figure S3C). To test whether the small subset of FRU-positive cells are involved in SMD, we expressed tetanus toxin light chain (*UAS[stop]TNT_active_)* with *Gr5a-* or *Gr66a-GAL4* drivers along with *fru^FLP^* to inhibit synaptic transmission in sexually dimorphic subsets of FRU-positive cells. Surprisingly (why is this surprising?), we found that SMD was abolished when *UAS-TNT* was expressed only in male-specific Gr5a-positive neurons (Figure 3D). As a control, we found that SMD was unaffected when we used each of these *GAL4* drivers in combination with *UAS[stop]TNT_inactive_* to express an inactive form of tetanus toxin light chain (Figure 3E). These data suggest that inhibition of synaptic transmission only in the male-specific Gr5a-positive neuronal population could specifically block the sensory inputs that induce SMD behavior.

Next, we tested for the effect of transforming these sensory neurons into the female form. It is well known that systemic expression of the female form of *tra* cDNA (*UAS-tra^F^*) in a male brain during development elicits female characteristics (Belote and Baker, 1987). Interestingly, we found that SMD was eliminated by the feminization of *Gr5a-GAL4* labeled cells, but not by expression of *UAS-tra^F^* in Gr66a-positive neuronal subsets (Figure 3F), suggesting that feminization of Gr5a-positive neurons nullify the male-specific sensory function of those cells to detect female cuticular pheromones and induce SMD behavior. All these data suggest that SMD requires the male-specific role of a subset of Gr5a-positive neurons. Together, these results suggest that the odorant binding proteins LUSH and coregulatory protein SNMP1 in male-specific Gr5a-positive neurons are crucial to send satiety signals to the central brain to produce SMD behavior.

### The circadian clock genes *Clock* and *cycle* are involved in SMD

One of the most crucial features of the central brain is to integrate temporal information with precise physiological responses. Evolution has favored different scales of biological timing from the 24 h circadian system to the seconds-to-minutes range of interval timing, and to the millisecond timing that is crucial for speech generation, motor control, sound localization, and consciousness of human being (Golombek, Bussi and Agostino, 2014). Among those different scales of biological timing, ‘interval timing’ is associated with essential behaviors such as foraging, learning, and decision making (Tucci, 2011). In human, it is the function of interval timing that allows us to experience the passage of real time subjectively. A human being can integrate action sequences, thoughts and behavior when interval timing functions properly. Our ability to detect emerging trends and to anticipate future outcomes also largely depends on the interval timing function of our brain (Block and Grondin, 2014; Buhusi and Meck, 2005). We suggested that the rival-induced extended mating, so called LMD behavior, could be a model for studying interval timing in *Drosophila melanogaster* (Kim, Jan and Jan, 2012, 2013). Also, here we suggest that SMD behavior can be an additional model system to investigate the neural basis of interval timing in the brain since both LMD and SMD behaviors involve decision-making procedures that take minutes.

The circadian clock is the most well understood features among biological timing in molecular terms (Hall, 2005). It constitutes of well-defined transcription/translation-based negative feedback loops that are controlled by core clock genes such as *timeless*, *period*, *Clock*, and *cycle* (Allada and Chung, 2010). In contrast to the circadian clock, the molecular components and the neural mechanisms for interval timing is not well understood. It has been suggested that the circadian clock may influence the interval timer, however, the molecular mechanisms is not known (Buhusi and Meck, 2005).

In *Drosophila*, the *Clock* (*Clk*)/*cycle* (*cyc*) dimer activates the transcription of *period* (*per*), *timeless* (*tim*), *vrille* (*vri*), *PAR domain protein 1* (*Pdp1*) and *clockwork orange* (*cwo*) genes, which in turn feedback to inhibit CLK-activated transcription or regulate *Clk* transcription (Zheng and Sehgal, 2008). In addition to regulating circadian rhythm of the animal, clock genes regulate the timing of non-circadian phenomena, such as the frequency of the male courtship song, developmental time, sleep length, cocaine sensitization, and giant fiber habituation (Hall, 2005). Our previous study revealed that *tim* and *per* are specifically involved in LMD (Kim, Jan and Jan, 2012).

To elucidate the relationship between circadian clock and interval timing, we tested the SMD behavior of core circadian clock gene mutants. We found that *Clk* and *cyc* mutant males did not show SMD whereas *per* and *tim* mutant males showed SMD (Figure 4A). Next, we asked whether the function of clock genes in the nervous system is sufficient to produce SMD behavior. Expression of CYC in *cyc* mutants with the pan-neuronal *GAL4* driver could rescue SMD behavior (Figure 4B), suggesting that neuronal expression of CYC protein is sufficient to elicit SMD behavior. Pigment-dispersing factor (PDF) expressing clock neurons are known to be the core pacemaker neurons in the fly brain (Kim, Jan and Jan, 2013). To test whether the function of the *cyc* gene in PDF-positive neurons are essential to elicit SMD behavior, we used the pan-neuronal *GAL4* driver combined with *pdf-GAL80* to block the function of *GAL4* specifically in PDF-positive cells. Interestingly, SMD defects in *cyc* mutants can be rescued with the introduction of CYC protein to neurons other than the PDF-expressing clock neurons (Figure 4B), suggesting that the function of CYC in PDF-expressing cells is dispensable for SMD behavior.

**Figure 4.**
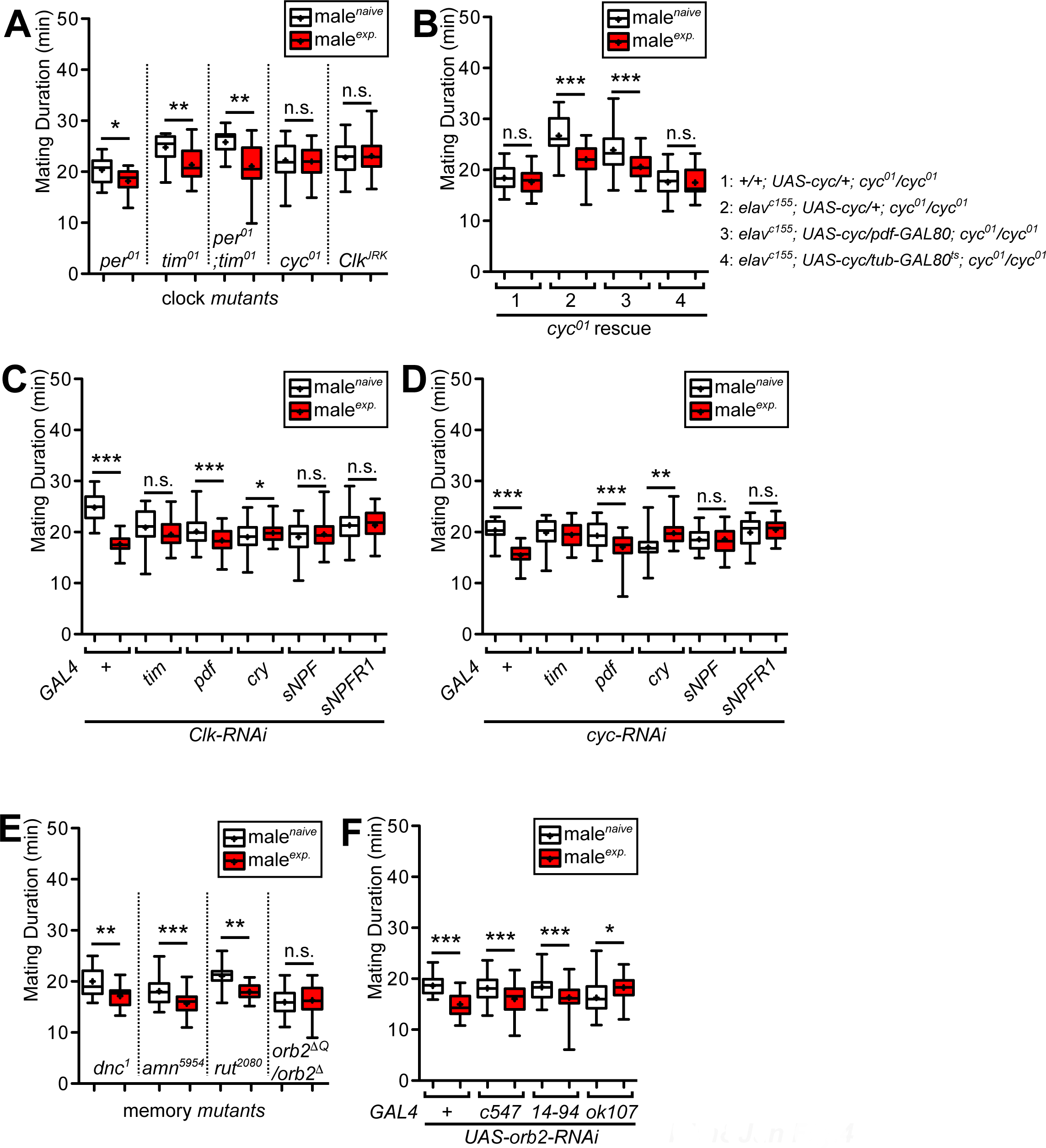
Circadian clock genes CLK and CYC are specifically involved with SMD. (A) MD assays of circadian clock mutants. The genotypes specified below the graphs. (B) MD assays of *cyc^01^* rescue experiments. Genotypes are indicated above the graphs. (C), (D) MD assay for *GAL4* mediated knockdown of CLK or CYC via *UAS-Clk-IR; UAS-dicer (Clk-RNAi) or UAS-cyc-IR; UAS-dicer (cyc-RNAi)*. Names of the *GAL4* drivers are indicated below the graphs. (E) MD assays of memory mutants. The genotypes specified below the graphs. (F) MD assay for *GAL4* mediated knockdown of ORB2 via *UAS-orb2-IR; UAS-dicer (orb2-RNAi)*. Names of the *GAL4* drivers are indicated below the graphs.

Next, we asked whether the function of clock genes on SMD behavior is adult-specific. Similar to previous findings that transgenic rescue of *cyc* mutants behavioral arrhythmia in adults depends on developmental *cyc* expression during metamorphosis (Goda, Mirowska, Currie, Kim, Rao, Bonilla and Wijnen, 2011), SMD could not be rescued by introducing CYC protein in an adult-specific manner indicating that the developmental function of CYC is required to elicit SMD behavior (Figure 4B). These data suggest that the developmental function of CLK/CYC in a subset of the neuronal population is required to induce SMD.

We then looked into the CLK/CYC function in SMD behavior. To identify the neuronal populations mediating the CLK/CYC function to elicit SMD, we screened for *GAL4* drivers that eliminate SMD behavior when combined with *Clk-RNAi* or *cyc-RNAi*. Knocking down expression of CLK or CYC in the majority of clock neurons eliminated SMD (*tim-GAL4* in Figure 4C and 4D). Expression of *Clk-RNAi or cyc-RNAi* in PDF-expressing lateral clock cells did not affect SMD (*pdf-GAL4* in Figure 4C and 4D). Removal of CLK or CYC from CRY-positive clock neurons eliminated SMD (*cry-GAL4* in Figure 4C and 4D), indicating that CLK/CYC expression in CRY-positive but PDF-negative cells is required for SMD, since CRY-positive neurons include all the PDF-positive cells in clock neurons. To confirm that PDF-positive neurons are dispensable for SMD behavior, we expressed *UAS-Kir2.1* in PDF-positive neurons in an adult specific manner to specifically inhibit the neuronal activity of those cells. We found that inactivation of PDF neurons did not affect SMD (*pdf* in Figure S4A). Taken together, these data suggest that CLK/CYC function in a subset of PDF-negative clock neurons is required to process sexual satiety information by female experience to produce SMD behavior.

### SMD requires memory circuitry residing in the mushroom and ellipsoid bodies

Males compete to find mates in a social environment (Wong and Candolin, 2005). In this circumstance, the ability of learning and memory can significantly increase the behavioral benefits of reproductive success of a male (Verzijden, ten Cate, Servedio, Kozak, Boughman and Svensson, 2012). In *Drosophila*, well-known behavioral paradigms such as “courtship conditioning” or “conditioned courtship suppression” have been widely used to study the theory of learning and the mechanisms of memory formation at the cellular and molecular level (Siegel and Hall, 1979). In most cases, the courtship conditioning paradigm has investigated the short-term memory (STM) lasting less than an hour. Long-term memory (LTM) forms in the fly brain when animals confront repetitive trials and a resting period, which lasts up to 24 h (Griffith and Ejima, 2009). Besides STM and LTM, mid-term memory (MTM) lasts from 1 to 3 hours. Moreover, two forms of long-term memory are distinguishable by training procedures, namely anesthesia-resistant memory (ARM) and long-term memory (LTM) (Heisenberg, 2003).

Sexual experiences can enhance the subsequent mating success through social learning and memory. Males can have a mating advantage over inexperienced males when they previously experienced mating. In this behavioral paradigm, courtship experience alone is not sufficient to provide a competitive advantage to the male. The copulation experiences play a significant role to reinforce the learning of males to perform more efficiently (Saleem, Ruggles, Abbott and Carney, 2014). Moreover, it has been reported that prior sexual experiences inhibit aggression between males. In this behavioral paradigm, 10 h or more experiences with females drastically suppress a male’s aggression against rival male. (Yuan, Song, Yang, Jan and Jan, 2014). Moreover, we previously reported that previous experience of rival-enriched environment is stored as long-term memory in ellipsoid body neurons through the function of memory genes, *rutabaga* (*rut*) and *amnesiac* (*amn*) (Kim, Jan and Jan, 2012).

Hence, we asked whether SMD requires memory circuits. The memory trace of sexual experiences for generating SMD behavior disappears between 12 h and 24 h (Figure S4B), suggesting that SMD requires some long-term memory trace. Next, we tested whether SMD was altered in classical learning and memory mutants (*dnc*, *rut*, *amn,* and *orb2)*. Interestingly, SMD was impaired only in *orb2* mutants, but not in *dnc, amn*, or *rut* mutants (Figure 4E). However, *orb2* mutants showed normal LMD behavior (Figure S4C). Thus *orb2* function is important for the memory formation involved in SMD. *Drosophila* Orb2 belongs to the CPEB2 subfamily, which functions to stimulate mRNA translation. In courtship conditioning, Orb2 is a crucial component to form a long-term memory (Keleman, Kruttner, Alenius and Dickson, 2007).

Genetic intervention has provided strong evidence that the mushroom body (MB) regions act as the seat of memory for odors (Heisenberg, 2003). In contrast, visual pattern memory in *D. melanogaster* is linked to the central complex, which includes the ellipsoid body (EB) and fan-shape body (FB) (Liu, Seiler, Wen, Zars, Ito, Wolf, Heisenberg and Liu, 2006; Pan, Zhou, Guo, Gong, Gong and Liu, 2009). To test which brain region is important for memory processing in SMD behavior, we used several *GAL4* lines that drive expression of *UAS-Kir2.1* in EB, MB, or FB brain regions. Expression of *UAS-Kir2.1* in the FB via *14-94*-*GAL4* had no effect on SMD (*14-94* in Figure S4A), whereas expressing *UAS-Kir2.1* in MB or EB regions abolished SMD (*ok107 and c547* in Figure S4A), suggesting that EB and MB are the brain regions to generate memory for SMD behavior.

Having found a requirement of the MB and EB in producing SMD behavior, we performed *GAL4*-mediated *RNAi* knockdown of *orb2* to identify the brain region where *orb2* functions to process memory involved in SMD behavior. Knocking down *orb2* expression in the MB disrupted SMD (*ok107* in Figure 4F), however, knocking down *orb2* in the EB or FB had no effect on SMD (*c547* and *14-94* in Figure 4F). These data suggest that the neural activity of MB and EB regions are both required for SMD, and *orb2* expression in MB is likely required to process the memory required for SMD.

### Neuropeptide sNPF is crucial component of SMD behavior

Neuropeptides regulate a wide range of animal behaviors. We found previously that LMD requires the functions of PDF and NPF but not sNPF in a subset of clock neurons (Kim, Jan and Jan, 2013). Since both LMD and SMD behaviors are related to modulated interval timing-based decision making, we tested whether the above neuropeptides expressed in clock neurons are involved with SMD behavior, by performing mating duration assays with mutants lacking a neuropeptide or its receptor. Since no *npf* mutant alleles are available (Nitabach and Taghert, 2008), we tested *npfR1* mutants (Burke, Huetteroth, Owald, Perisse, Krashes, Das, Gohl, Silies, Certel and Waddell, 2012; Krashes, DasGupta, Vreede, White, Armstrong and Waddell, 2009). Among the mutants tested, males of mutant for *npfR1* or *pdf* but not *sNPF* (Lee, Kwon, Lee, Kwon, Min, Jung, Kim, You, Tatar and Yu, 2008b) displayed SMD behavior (Figure 5A and Figure S5A), indicating that SMD behavior only requires the function of neuropeptide sNPF, but not PDF or NPF.

**Figure 5.**
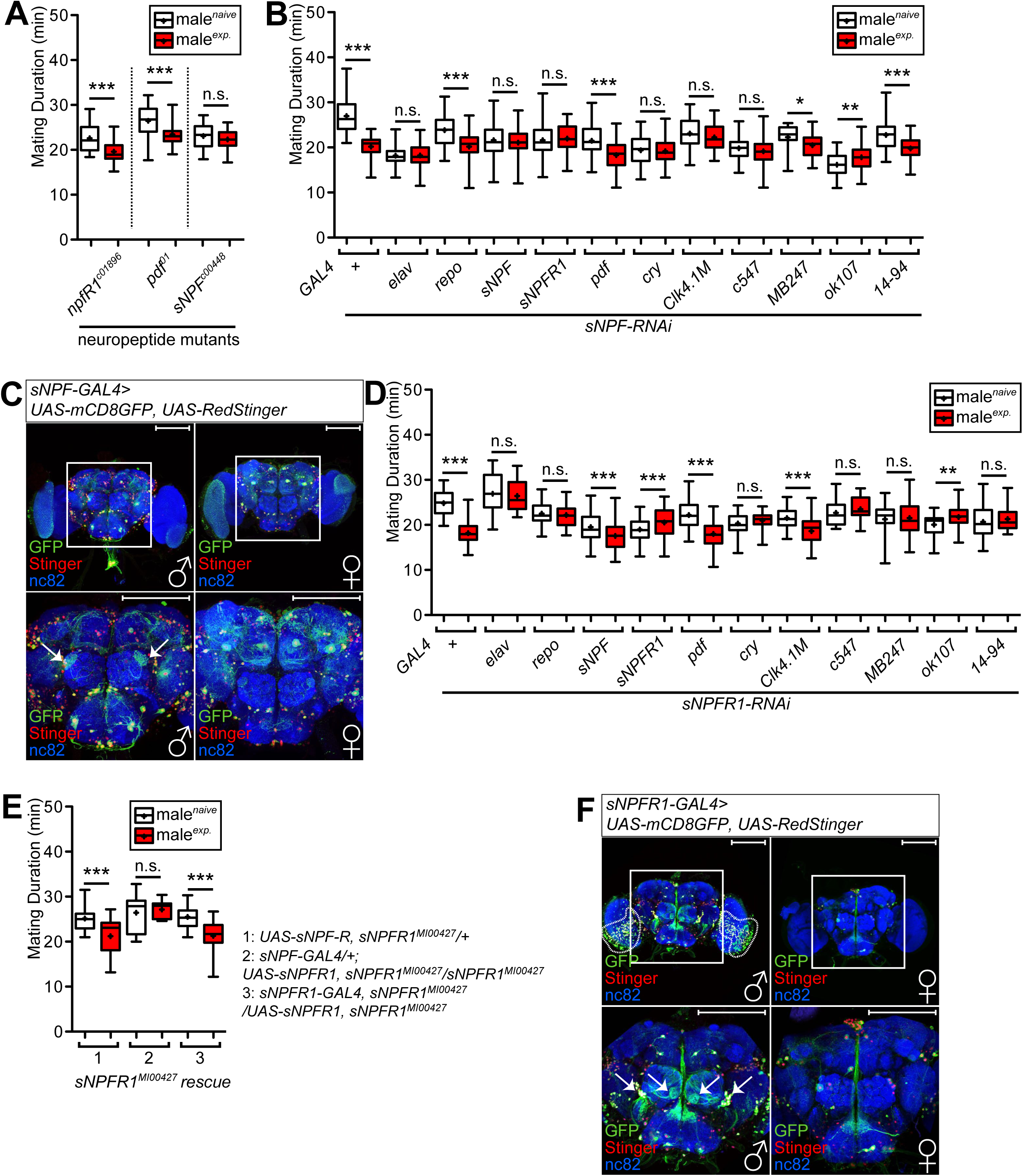
sNPF signaling is required to produce SMD. (A) MD assays of neuropeptides and neuropeptides receptor mutants. The genotypes specified below the graphs. (B) MD assays for *GAL4* mediated knockdown of sNPF via *UAS-sNPF-IR; UAS-dicer (sNPF-RNAi)*. Names of the *GAL4* drivers are indicated below the graphs. (C) Brains of flies expressing *sNPF-GAL4* together with *UAS-mCD8GFP; UAS-RedStinger* were immunostained with anti-GFP (green), anti-DsRed (red) and nc82 (blue) antibodies. White boxes indicate the magnified regions of interest presented next bottom panels. Scale bars represent 100 μm. (D) MD assays for *GAL4* mediated knockdown of sNPF-R1 via *UAS-sNPF-R-IR; UAS-dicer (sNPF-R-RNAi)*. Names of the *GAL4* drivers are indicated below the graphs. (E) MD assays of *sNPF-R^MI004271^* rescue experiments. Genotypes are indicated below the graphs. (F) Brains of flies expressing *sNPF-R-GAL4* together with *UAS-mCD8GFP; UAS-RedStinger* were immunostained with anti-GFP (green), anti-DsRed (red) and nc82 (blue) antibodies. White boxes indicate the magnified regions of interest presented next bottom panels. Scale bars represent 100 μm.

Insect neuropeptides are expressed in discrete stereotypic neuronal populations in the central nervous system. Short neuropeptide F (sNPF) is the *Drosophila* homologue of the mammalian neuropeptide Y (NPY), which controls food consumption. sNPF is expressed in the nervous system and regulates food intake and body size (Lee et al., 2008b). sNPF is widely expressed in a variety of neurons in the fly brain including Kenyon cells of mushroom bodies. There are several thousands of diverse types of neurons in the adult fly brain expressing *snpf* transcript and sNPF peptide. It is known that the most of these neurons are inherent interneurons of mushroom bodies. Also, sNPF is expressed in many interneurons of the CNS, olfactory receptor neurons (ORNs) located in antennae, and the subpopulation of neurosecretory cells innervating the corpora cardiac and aorta (Nassel, Enell, Santos, Wegener and Johard, 2008). Even though the names of the two peptide genes (NPF and sNPF) are similar, previous reports showed that they play roles in two functionally distinct signaling pathways (Johard, Enell, Gustafsson, Trifilieff, Veenstra and Nassel, 2008; Lee, Kwon, Lee, Kwon, Min, Jung, Kim, You, Tatar and Yu, 2008a; Lee, You, Choo, Han and Yu, 2004; Wu, Wen, Lee, Park, Cai and Shen, 2003). These reports are consistent with our result. Our data suggest that the functions of these two neuropeptides do not overlap since sNPF but not NPF is involved with SMD behavior (Figure 5A), whereas NPF but not sNPF is involved with LMD (Kim, Jan and Jan, 2013).

We next wanted to identify the neurons that express sNPF to mediate SMD behavior. We used a previously verified *UAS-sNPF-RNAi* line (Lee, Kwon, Lee, Kwon, Min, Jung, Kim, You, Tatar and Yu, 2008b) in combination with various *GAL4* drivers to knock down sNPF expression in discrete populations of *GAL4* expressing cells. The brain regions labeled by the *GAL4* drivers used in this study have been identified previously (Kim, Jan and Jan, 2012, 2013), as diagramed in Figure S5B. We found that SMD was abolished by expression of *sNPF-RNAi* in all neuronal population (via pan-neuronal *elav-GAL4* driver, *elav* in Figure 5B) but not in glial cells (via *repo-GAL4*, *repo* in Figure 5B). We then tested the function of sNPF expression in sNPF-expressing cells and in cells expressing its receptor sNPFR1 using previously verified sNPF-GAL4 and sNPFR1-GAL4 strains (Lee, Kwon, Lee, Kwon, Min, Jung, Kim, You, Tatar and Yu, 2008a). Knock down of sNPF using *sNPF-* or *sNPFR1-GAL4* abolished SMD (*sNPF* and *sNPFR1* in Figure 5B), indicating that sNPF-expression in a subset of cells that also express its receptor are important to induce SMD.

As described above, the circadian clock genes *Clk* and *cyc* are intimately involved with SMD behavior (Figure 4). Therefore, we tested *GAL4* drivers that label different populations of clock neurons to identify the subset of clock neurons required for sNPF signaling. Interestingly, SMD remained intact with expression of *sNPF-RNAi* in PDF-expressing neurons (*pdf* in Figure 5B), but was abolished by expression of *sNPF-RNAi* in CRY-positive cells, which include most of the lateral neurons and a small subset of dorsal neurons (*cry* in Figure 5B). Thus, sNPF expression in neurons that express CRY but not PDF is required for SMD behavior. Expression of *sNPF-RNAi* in a subset of dorsal neurons (*Clk4.1M* in Figure 5B) eliminated SMD, indicating that sNPF signaling in a subset of CLK-positive neurons is required for SMD.

We showed that SMD behavior requires memory circuit located in mushroom bodies (MB) and ellipsoid bodies (EB) (Figure S4A). Therefore, we next asked if sNPF signaling is critical to memory formation and processing for SMD behavior. We reasoned that knock down of sNPF signaling in mushroom bodies or ellipsoid bodies would disrupt SMD behavior if sNPF signaling plays a pivotal role in memory formation/processing. To characterize the relationships between sNPF signaling and memory circuits in SMD, we used different *GAL4* drivers to express *sNPF-RNAi* in ellipsoid bodies (*c547-GAL4*), mushroom bodies (*MB247-* and *ok107-GAL4*), and fan-shape bodies (FB) (*14-94-GAL4*). Whereas the expression of sNPF in EB neurons is required for SMD (*c547* in Figure 5B), sNPF signaling in FB is dispensable for SMD (*14-94* in Figure 5B). When sNPF expression is reduced in MB via *ok107-GAL4*, SMD was eliminated, however knocking down sNPF expression via *MB247-GAL4*, which labels more restricted regions of the MB, did not affect SMD behavior (*ok107* and *MB247* in Figure 5B), indicating that sNPF signaling in MB neurons labeled by *ok107-GAL4* but not *MB247-GAL4* is crucial for SMD behavior (Jenett, Schindelin and Heisenberg, 2006).

We next asked what happens if an excessive amount of sNPF exists in a subpopulation of neurons that normally expresses sNPF or its receptor sNPFR1. Overexpression of *UAS-sNPF* via *sNPF-GAL4* or *sNPFR1-GAL4* had no effect on SMD behavior (Figure S5C), indicating that increasing the amount of this neuropeptide in sNPF-positive or sNPFR1-positive neurons has no detrimental effect on SMD behavior.

Previous studies have shown that sNPF and its receptor sNPFR1 modulate feeding behavior and growth in fruit fly (Lee, Kwon, Lee, Kwon, Min, Jung, Kim, You, Tatar and Yu, 2008b; Lee, You, Choo, Han and Yu, 2004). However, those studies investigated the role of sNPF signaling by widespread overexpression. Therefore, sNPF action in specific circuits has not been addressed. To identify neurons that must express sNPF to mediate SMD, we first characterized the expression pattern of the *sNPF-GAL4* driver using the reporter *UAS-mCD8GFP* (a cell membrane marker) together with *UAS-RedStinger* (a nuclear marker) (Figure 5C), or *UAS-Denmark* (a dendritic marker) together with *UAS-sytGFP* (a presynaptic marker) (Figure S5D and S5E) (Nicolai, Ramaekers, Raemaekers, Drozdzecki, Mauss, Yan, Landgraf, Annaert and Hassan, 2010). This sNPF-GAL4 driver generates an expression pattern that recapitulates the pattern revealed by sNPF antibody immunocytochemistry (Nässel, Enell, Santos, Wegener and Johard, 2008). We identified more than 200 cells that were labeled by the *sNPF-GAL4* driver, including MB neurons, s-LNv (white arrows in Figure S5E), a subset of PI (*pars intercerebrails*) neurons, neurons in the SOG (suboesophageal ganglion), and a subset of antenna lobe neurons specifically labeled in the male fly brain (white arrows in Figure 5C), indicating that some sNPF expressing cells are sexually dimorphic. We cannot exclude the possibility that some small sNPF-GAL4 labeled cells may have escaped detection.

Next we asked how sNPFR1, the only-known *Drosophila* sNPF receptor (NPFR76F; CG7395; sNPFR1) (Feng, Reale, Chatwin, Kennedy, Venard, Ericsson, Yu, Evans and Hall, 2003b; Mertens, Meeusen, Huybrechts, De Loof and Schoofs, 2002), mediates sNPF signaling to modulate SMD behavior. We used *sNPFR1-RNAi* combined with various *GAL4* drivers to identify the neurons that express the sNPF receptor to mediate SMD behavior (Figure 5D). Expression of *sNPFR1-RNAi* in neuronal or glial populations eliminated SMD (*elav* and *repo* in Figure 5D), indicating that glial cells expressing sNPFR1 are also required for SMD behavior. SMD was intact when sNPFR1 expression was reduced in sNPF expressing cells via *sNPF-GAL4* (*sNPF* in Figure 5D), indicating that sNPFR1 (receptor) expression among sNPF-positive cells, potentially for autocrine signaling, is not required for SMD behavior.

Next we tested *GAL4* drivers that label different populations of clock neurons to identify the subset of clock neurons required for sNPFR1 signaling. Although SMD remained intact with expression of *sNPFR1-RNAi* in PDF-expressing neurons (*pdf* in Figure 5D), SMD was abolished by expression of *sNPFR1-RNAi* in CRY-positive cells (*cry* in Figure 5D). Thus, sNPFR1 expression in neurons that express CRY but not PDF is required for SMD. Expression of *sNPFR1-RNAi* in a subset of dorsal neurons (*Clk4.1M* in Figure 5D) did not eliminate SMD. This result implies that sNPFR1 expression is not required in the CLK-positive small subset of dorsal neurons to induce SMD behavior, even though sNPF expression in a subset of CLK-positive neurons is required for SMD (*Clk4.1M* in Figure 5B). Reducing sNPFR1 signaling in MB, EB, and FB neurons eliminated SMD (*c547*, *MB247*, *ok107*, and *14-94* in Figure 5D), indicating that sNPFR1 signaling in these memory circuits are required to elicit SMD.

To delineate the relationship between sNPF signaling and its known receptor sNPFR1, we performed rescue experiments using *sNPFR1^MI00427^* mutants which are homozygous-lethal. As expected, SMD was restored in *sNPFR1^MI00427^* mutants by expressing the *UAS-sNPFR1* transgene via the *sNPFR1-GAL4* driver (lane 3 in Figure 5E). Interestingly, expression of the *UAS-sNPFR1* transgene via the *sNPF-GAL4* driver in *sNPFR1^MI00427^* mutants rescued the lethality but not SMD behavior (lane 2 in Figure 5E). These studies reveal that sNPFR1 expression in sNPF-positive cells is crucial for survival, but not sufficient to fully elicit SMD behavior, indicating that sNPFR1 expression in regions other than sNPF-expressing cells are required to elicit intact SMD behavior.

To identify neurons that express sNPFR1 to mediate the SMD behavior, we characterized the expression pattern of the *sNPFR1-GAL4* driver using the reporter *UAS-mCD8GFP* together with *UAS-RedStinger* (Figure 5F). We found more than 100 cells labeled by the *sNPFR1-GAL4* driver including MB neurons and a subset of PI neurons. A subset of cells in the antenna lobe and cells located in the ventral optic lobe (white arrows in Figure 5F) were labeled only in male fly brains, indicating that a portion of sNPFR1 expressing cells are sexually dimorphic.

Since we already determined that Gr5a-positive sensory neurons are required for SMD (Figure 2 and 3), we next asked whether sNPF signaling in these sensory cells are also required for SMD. Previous studies have implicated sNPF signaling in gustatory sensory transduction (Ci, Wu and Su, 2014; Inagaki, Panse and Anderson, 2014). Expression of *sNPF-RNAi* in Gr5a-positive, but not Gr66a-positive cells, abolished SMD (Figure S5F), indicating that sNPF signaling in Gr5a-positive neurons is required to elicit SMD behavior. In addition, we found that the function of CLK/CYC in neurons mediating sNPF signaling is crucial for SMD (*sNPF* and *sNPFRR1* in Figure 4C and D), indicating that sNPF signaling is connected to gustatory neural circuits and circadian clock neurons for modulating SMD behavior.

### Sexually dimorphic sNPF-expressing neurons are required for SMD

Sexual dimorphism refers “the differences in appearance between males and females of the same species, such as color, shape, size, and structure, that are caused by the inheritance of one of the other sexual pattern in the genetic material”. In *Drosophila*, the neural sex determination gene, *fruitless* (*fru*) has been identified and investigated. FRU is translated only in males to regulate the transcription of a set of downstream effectors. At least in neuronal populations, expression of FRU in female induces male-like behaviors (Yamamoto, 2007). We previously showed that FRU-positive, sexually dimorphic neuropeptide F (NPF) expressing neurons modulate LMD behavior (Kim, Jan and Jan, 2013).

As we have shown above, *sNPF-* and *sNPFR1-GAL4* expression patterns exhibit sexual dimorphism (Figures 5C and 5F). We therefore wanted to identify the sexually dimorphic neurons among sNPF-positive cells. First, we used *sNPF-GAL4* to identify sexually dimorphic cells using intersectional method with *fru^FLP^* as described above (Figure 3B). Interestingly, we found ~40 FRU-positive and sNPF-positive neurons in the male brain (*sNPF-GAL4* in Figure 6A). When *sNPFR1-GAL4* was used to identify sexually dimorphic cells, strong GFP fluorescence near the antenna lobe was detected in the male brain (*sNPFR1-GAL4* in Figure 6A). These data suggest the presence of FRU-positive cells among the sNPF- and sNPFR1-postive neuronal populations.

**Figure 6.**
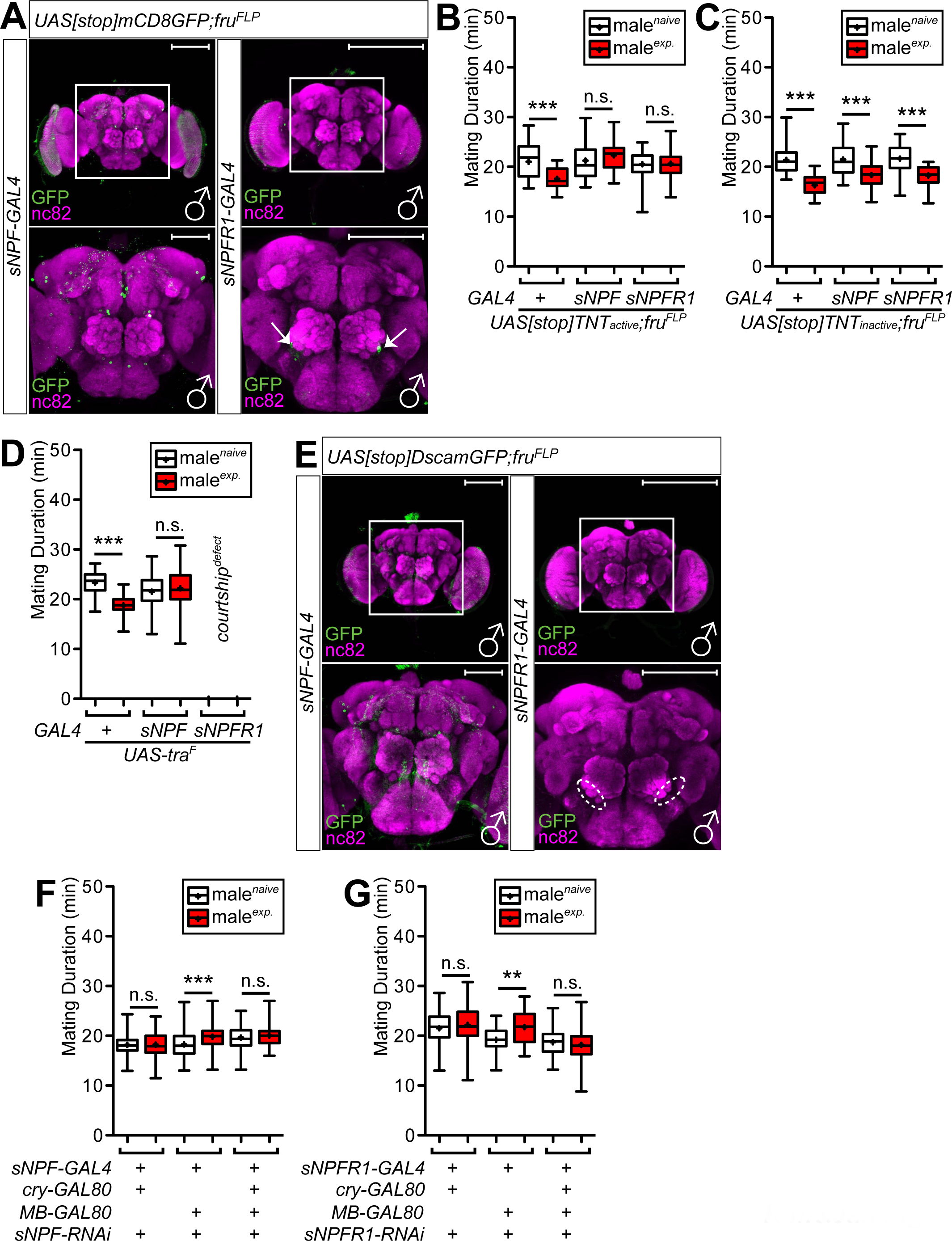
Both male-specific and redundant sNPF signaling modulates SMD. (A) Brains of flies expressing *sNPF-GAL4 or sNPF-R-GAL4* together with *UAS[stop]mCD8GFP; fru^FLP^* were immunostained with anti-GFP (green) and nc82 (magenta) antibodies. White boxes indicate the magnified regions of interest presented the bottom panels. Scale bars represent 100 μm. (B) MD assays of *sNPF-* and *sNPF-R-GAL4* drivers for inactivation of synaptic transmission of male-specific neurons among each *GAL4*-labeled neurons via *UAS[stop]TNTactive; fru^FLP^*. (C) Control experiments of **Figure 6B** with *inactive form of UAS-TNT using UAS[stop]TNTinactive; fru^FLP^*. (D) MD assays for *sNPF-* and *sNPF-R-GAL4* drivers for feminization of neurons via *UAS-tra^F^*. Since feminizing *sNPF-R-GAL4* labeld neurons seriously affect courtship activity, MD assays could not be done. (E) Brains of flies expressing *sNPF-GAL4 or sNPF-R-GAL4* together with *UAS[stop]DscamGFP; fru^FLP^* were immunostained with anti-GFP (green) and nc82 (magenta) antibodies. White boxes indicate the magnified regions of interest presented the bottom panels. Dashed circles in represent the clearly disappeared GFP signals compared to **Figure 6A** (white arrows). Scale bars represent 100 μm. (F) and (G) MD assays for *GAL4/GAL80* mediated knockdown of sNPF or sNPF-R1 via *UAS-sNPF-IR; UAS-dicer (sNPF-RNAi) or UAS-sNPF-R-IR; UAS-dicer (sNPF-R-RNAi)*. Names of the *GAL4/GAL80* drivers’ combination are indicated below the graphs.

We then examined the function of these FRU-positive neurons on SMD behavior. To test whether the small subset of neurons positive for both *fru* and *sNPF* or positive for both *fru* and *sNPFR1*, are involved in SMD, we expressed tetanus toxin light chain (*UAS[stop]TNT_active_)* with *sNPF-* or *sNPFR1-GAL4* drivers along with *fru^FLP^* to inhibit synaptic transmission in sexually dimorphic subsets of *fru*-positive cells. Expression of *UAS-TNT* in *fru-* and *sNPF*-positive, or *fru-* and *sNPFR1-*positive, neurons, abolished SMD behavior (Figure 6B). As a control, we found that SMD was unaffected when we used each of these *GAL4* drivers in combination with *UAS[stop]TNT_inactive_* to express an inactive form of tetanus toxin light chain (Figure 6C). Similar results (Figure S6G) were obtained using the cell autonomous toxin *Ricin A (UAS[stop]RicinA*), which encodes only the catalytic subunit of the toxin that can enter the cell to cause cell death (Hidalgo, Urban and Brand, 1995). These data suggest that those FRU-positive neurons that are involved in sNPF signaling are crucial to elicit SMD behavior.

Next, we asked if only the masculinized form of those neurons can induce male-specific SMD behavior. Feminization of *sNPF-GAL4* labeled neurons using *UAS-tra^F^* expression eliminated SMD, suggesting that the male-specific function of s*NPF*-positive neurons is required to elicit SMD (*sNPF* in Figure 6D). Male flies did not show any significant courtship activity when *sNPFR1*-GAL4 labeled cells were feminized. These males failed to compete for a mate, and therefore we could not measure mating duration in these flies (*sNPFR1* in Figure 6D). These results suggest that the male-specific function of a subset of *sNPFR1*-positive cells is critical to induce courtship behavior. In addition, we noticed that the expression of GFP fluorescence in SOG regions was lost when sNPFR1-positive cells were feminized via *UAS-tra^F^* expression (compare Figure 5F with Figure S6F), raising the possibility that a subset of sNPFR1-positive cells that function in SOG regions is responsible for normal courtship behavior.

Next we used the dendritic marker *UAS[stop]DscamGFP* to label neurons positive for both *fru* and *sNPF,* or positive for both *fru* and *sNPFR1*. We found large dendritic arbors spanning from dorsal to SOG regions as revealed by using the *sNPF-GAL4* driver (*sNPF* in Figure 6E). No GFP expressing processes positive for both *fru* and *sNPFR1* could be found with the dendritic marker (Figure 6E), suggesting that the GFP fluorescence shown in Figure 6A corresponds to presynaptic terminals rather than dendrites.

As described above, both *sNPF-GAL4* and *sNPFR1-GAL4* label thousands of cells in the fly brain. To further narrow down the population of sNPF- *and* sNPFR1-expressing neurons involved in SMD, we used different *GAL80* lines to block the function of *sNPF-* or *sNPFR1-GAL4* drivers in brain regions with *GAL80* expression. With these *GAL4/GAL80* combinations, we knocked down *sNPF or sNPFR1* expression using *sNPF-RNAi* or *sNPFR1-RNAi* respectively (Figure 6F and 6G). SMD was abolished when sNPF was knocked down using *sNPF-GAL4* with *cry-GAL80* that expresses GAL80 in CRY-positive neurons, with *MB-GAL80* that expresses specifically in the mushroom body, or with both *cry-GAL80* and *MB-GAL80* (Figure 6F). This indicates that the function of sNPF signaling in CRY- and MB-positive neurons is not sufficient to fully induce SMD. Similar results were obtained when sNPFR1 was knocked down using *sNPFR1-GAL4* with *cry-GAL80* or *MB-GAL80* (Figure 6G). We tested the expression of these *GAL4/GAL80* combinations using *UAS-mCD8GFP and UAS-RedStinger,* confirming the suppression of GFP expression via *GAL80* expression (Figures S6A-E). All these data suggest that sNPF/sNPFR1 signaling in multiple brain regions is required to generate SMD.

### Sexual experience affects the properties of synapses and activity of neurons involved in sNPF signaling

The most probable place which reflects plastic changes in neurons are synapses, the junction between neurons. Various studies have revealed that the synaptic terminals of sensory and interneurons are pivotal sites of neuromodulation (Sweatt, 2016). In *Drosophila*, previous studies have shown that starvation increase the sugar sensitivity of gustatory neurons leading to altered behavioral responses to sugar (Gaudry and Kristan, 2009), which may involve the neuropeptide sNPF signaling that functions locally in olfactory tissues to modulate this neuronal plasticity (Root, Ko, Jafari and Wang, 2011). In this scenario, sNPF signaling can create internal states of starved or well-fed flies so that sensory inputs should be gated and modulated by those internal states (Kennedy, Asahina, Hoopfer, Inagaki, Jung, Lee, Remedios and Anderson, 2014).

As shown above, we have demonstrated that SMD behavior displays behavioral plasticity, which indicates that males reversibly change the mating duration dependent on their environmental context (Figures 1 and S1). Thus we asked if we could find neuronal populations presenting synaptic plasticity dependent on male’s sexual experiences. To selectively label nerve terminals, we used neuronal synaptobrevin-GFP chimera (*nsybGFP*), which allows visualizing the terminals of axonal projections. The expression of *nsybGFP* alters neither morphology nor circuitry of neurons thus it could be used as a tracer of axonal projections of neural circuits in the fly brain (Vosshall, Wong and Axel, 2000). Many studies have used *nsybGFP* to measure activity-dependent chronic plasticity in olfactory circuits (Sachse, Rueckert, Keller, Okada, Tanaka, Ito and Vosshall, 2007) (Devaud, Acebes and Ferrus, 2001). Further, GFP fluorescence of *nsybGFP* allows visualization of synapse formation, retraction, and maturation during embryonic development or metamorphosis, and for determining projection patterns and target sites of CNS neurons in the adult brain (Estes, Ho, Narayanan and Ramaswami, 2000).

To look into the synaptic plasticity of the sNPF-expressing neuronal population, we used *UAS[stop]nSybGFP* with *fru^FLP^* (Yu, Kanai, Demir, Jefferis and Dickson, 2010b) in combination with the *sNPF-GAL4* driver to analyze the minimal neural populations functionally responsible for SMD behavior (Figure 6A), given that the *sNPF-GAL4* driver labels numerous cells in the adult brain (compare Figure S7A with Figure 7A). Interestingly, the number of *nSybGFP* particles increased in experienced males (Figure 7B), while the average size of those particles was similar between naïve and experienced males (Figure 7C). The percent of area covered by *nsybGFP* particles increased in the brains from experienced males (Figure 7D). Interestingly, there was a large increase in the number and intensity of *nsybGFP* particles near SOG regions in experienced males (white circles in Figure 7A), indicating that gustatory inputs from female partners enhance sNPF signaling within specific CNS regions. We also used *UAS[stop]DscamGFP* with *fru^FLP^* in combination with *sNPF-GAL4* to analyze how sexual experience modulates the dendritic morphology of responsible neurons (Figure S7B). The number of *DscamGFP* particles (Figure S7C), average size (Figure S7D), and percent of area (Figure S7E) were all increased in experienced males, indicating that chronic sexual experience modulates the dendritic morphology of sexually dimorphic sNPF-positive neurons responsible for inducing SMD behavior. We suggest that this neuronal reorganization in sNPF-expressing neurons alters the internal state of the male brain that enables the male’s decision to terminate mating earlier than before.

**Figure 7.**
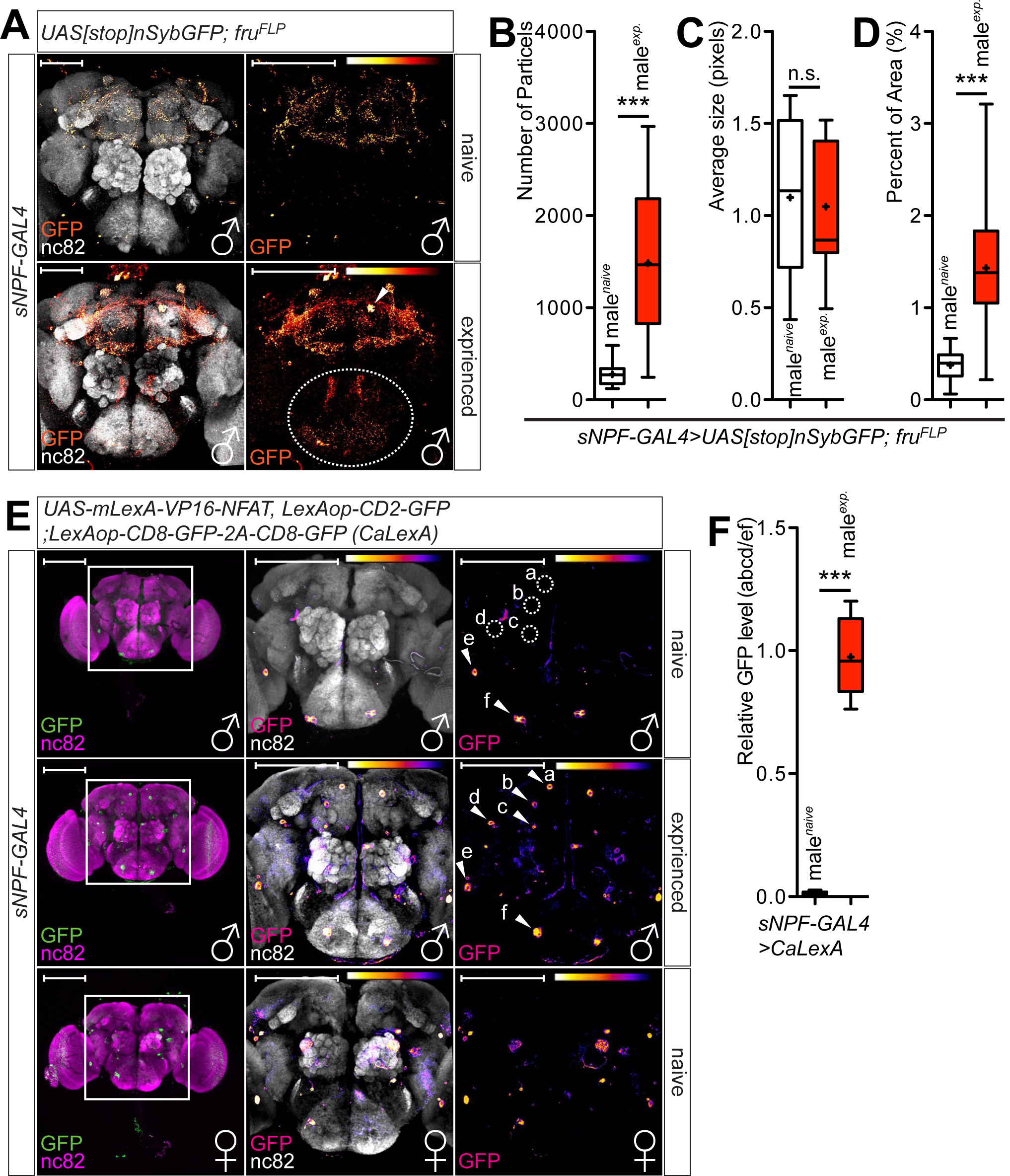
Sexual experience alters the number of synapses and the neural activity of neuropeptide expressing neurons. (A) Flies expressing *sNPF-GAL4* with *UAS[stop]n-sybGFP; fru^FLP^* were naively reared (upper panels) or experienced with virgin females 1 day before dissection (bottom panels). Brains of flies were immunostained with anti-GFP (red hot) and nc82 (grey) antibodies. GFP is pseudocolored “red hot”. Dashed circles represent the clearly increased GFP signals compared to upper panel. Scale bars represent 100 μm. (B) Number of particles shown in (A) was quantified using *ImageJ* software. (C) Average size of particles shown in (A) was quantified. (D) Percent of brain area covered by particles shown in (A) are quantified. See EXPERIMENTAL PROCEDURES for details. (E) Brains of naïve male (top), experienced male (middle), and female (bottom) flies expressing *sNPF-GAL4* along with *LexAop-CD2-GFP; UAS-mLexA-VP16-NFAT, LexAop-CD8-GFP-2A-CD8-GFP* were dissected then immunostained with anti-GFP (green) and anti-nc82 (magenta) antibodies. GFP is pseudocolored “fire” in the middle and the right panels. Warmer colors reflect increased signal intensity. Cells showing strong GFP signals were labeled “a-f” from the top to bottom. White arrows indicate the subset of neurons showing strong GFP signals; dashed circles indicate the subset of neurons showing reduced GFP signals compared to the different rearing condition. (F) GFP fluorescence of “a-d” regions is normalized by “e-f” region. See **EXPERIMENTAL PROCEDURES** for detailed quantification methods.

Internal states of experienced male may be correlated with the number of synapses and morphology of dendrites. To find out whether neuronal activity is altered in neurons involved in SMD, we used the CaLexA (calcium-dependent nuclear import of LexA) system (Masuyama, Zhang, Rao and Wang, 2012), to measure neuronal activity based on the activity dependent nuclear import of a nuclear factor of activated T cells (NFAT). Since SMD requires chronic exposure to females for at least 12-24 h, the repeated sensory inputs could conceivably result in the accumulation of the engineered transcription factor in the nuclei of active neurons *in vivo*. This system has been successfully applied to measure Ca^2+^ levels over the course of multiple hours inside the brain (Guo, Yu, Jung, Abruzzi, Luo, Griffith and Rosbash, 2016; Kayser, Yue and Sehgal, 2014b; Koh, He, Gorur-Shandilya, Menuz, Larter, Stewart and Carlson, 2014; Liu, Liu, Tabuchi and Wu, 2016; Park, Dus, Kim, Abu, Kanai, Rudy and Suh, 2016).

We first examined sNPF-expressing neurons, since they are a critical modulator of SMD behavior. As shown above, sexual experiences substantially alter the synaptic terminals and dendritic morphology of sNPF-positive neurons (Figure 7A and Figures S7A-B). Indeed, the neuronal activity of some *sNPF-GAL4* labeled neurons was altered by sexual experience. Male flies harboring *sNPF-GAL4* and *LexAop-CD2-GFP; UAS-mLexA-VP16-NFAT, LexAop-CD8-GFP-2A-CD8-GFP (CaLexA)* showed robust fluorescence in three dorsomedial neurons and one dorsolateral neuron after 1 day of sexual experience (labeled as a-d in Figure 7E). In contrast, one neuron in the lateral and one neuron in the ventromedial regions showed constant robust fluorescence in both naïve and experience males (labeled as e-f in Figure 7E). Thus, sexual experience increased the fluorescence signal of dorsal neurons compared to that of ventral neurons (abcd/ef fluorescence) in males (Figures 7F). GFP fluorescence in female brains showed different patterns, suggesting that it is the male-specific sNPF-positive neurons that are activated after sexual experience (bottom panels in Figure 7E). Male flies harboring *sNPFR1-GAL4* and *CaLexA* showed diminished GFP fluorescence in ventromedial antennal lobe neurons, as compared to dorsomedial antennal lobe neurons, after 1 day of sexual experience (labeled as a-b in Figure S7F). Thus sexual experience decreased the fluorescence signal of ventromedial antennal lobe neurons compared to that of dorsomedial antennal lobe neurons (b/a fluorescence) in males (Figures S7G). As control, we examined PDF- and NPF-expressing neurons that are not involved in SMD behavior by using the *pdf-GAL4* or *npf-GAL4* driver to assess their neuronal activity. We did not find any significant changes in GFP fluorescence between PDF- and NPF-expressing neurons by sexual experience (Figures S7H-J), indicating that sNPF signaling but not PDF or NPF signaling is specifically involved in generating SMD behavior via sexual experience.

## DISCUSSION

Our study provides evidences that male flies invest less time for copulation when they are sexually satiated. Males retain a memory of sexual experience for several hours and economize mating duration accordingly. This behavior relies primarily on the gustatory input (Figure 2) that originates from female pheromones produced by oenocytes (Figure 1), indicating that contact chemoreception is required for SMD induction. Male-specific Gr5a-positive cells are sensory neurons required for recognizing the presence of females and these gustatory neurons are connected to the central brain region to process information for SMD behavior (Figure 3). We further show that the circadian clock genes *Clk* and *cyc,* but not *per* or *tim,* are important for SMD behavior. In addition, we found that the function of CLK/CYC in a subset of clock neurons within the circadian rhythm circuit is important for SMD behavior (Figure 4A-D). SMD requires the function of ellipsoid bodies and mushroom bodies within the CNS for memory processing, and this processing depends on *orb2* function within the mushroom bodies (Figure 4E-F). We provide evidence for the crucial involvement of the neuropeptide sNPF, but not PDF or NPF, in the modulation of reproductive behavior by the male’s prior sexual experience (Figure 5). Sexually dimorphic sNPF/sNPFR1-positive neurons are necessary to generate SMD (Figure 6). Remarkably, sexual experience alters the synaptic structure and the neuronal activity of a subset of neurons expressing sNPF likely reflecting neuronal modulation relevant for the SMD behavior (Figure 7).

### How does sNPF signaling orchestrate multiple functions of brain dynamics?

Neuropeptides can function in a diverse way. In the central brain, they act as neuromodulators or cotransmitters. Neuropeptides also can play a role as circulating hormones via the hemolymph and as neuromediators when released by neurons to influence peripheral targets (Nassel and Winther, 2010). Neuropeptide sNPF is widely distributed in various types of neurons in the brain and VNC of the fly. It has been suggested that sNPF is a multifunctional neuropeptide (Nassel, Enell, Santos, Wegener and Johard, 2008). For example, the effect of sNPF on growth may be limited to a restricted subset of the sNPF-expressing neurons near insulin-producing cells in pars cerebrails (PI). Thus the regulation of growth by sNPF signaling is only a part of a spectrum of sNPF functions in physiology and behavior in *Drosophila* (Lee, Kwon, Lee, Kwon, Min, Jung, Kim, You, Tatar and Yu, 2008a).

The multiple functions of sNPF in fruit fly have been suggested as follows: It plays a role in olfaction at the synapses between olfactory receptor neurons (ORN) and second order neurons in the antennal lobe. It functions in the mushroom bodies to process olfactory information and modulate olfactory learning to support the role of mushroom bodies. It also functions as a neuromodulator in the central complex of the fly brain to control locomotion. As described above, the upstream signal of the insulin-producing cells also functions as circulating neurohormones. Finally, it likely acts as modulator/cotransmitter functions in various brain circuits and the ventral nerve cord (Johard, Enell, Gustafsson, Trifilieff, Veenstra and Nassel, 2008). To understand the functional diversity of sNPF signaling in regulating various functions of brain dynamics, we need to interfere with the function of sNPF more precisely in discrete circuits. In this context, one might expect the various function of sNPF can be achieved by modulating its receptor expression in diverse neuronal types. There is only a single *Drosophila* sNPF receptor identified so far (NPFR76F; CG7395; sNPFR1) (Feng, Reale, Chatwin, Kennedy, Venard, Ericsson, Yu, Evans and Hall, 2003a; Mertens, Meeusen, Huybrechts, De Loof and Schoofs, 2002; Reale, Chatwin and Evans, 2004). It already has been postulated that this receptor can segregate the function of sNPF by coupling it to different signaling pathways (Feng, Reale, Chatwin, Kennedy, Venard, Ericsson, Yu, Evans and Hall, 2003a; Lee, Kwon, Lee, Kwon, Min, Jung, Kim, You, Tatar and Yu, 2008a; Mertens, Meeusen, Huybrechts, De Loof and Schoofs, 2002; Reale, Chatwin and Evans, 2004).

Consistent with previous reports, our study provides a new line of evidence that a small number of the neuronal population requires the function of sNPF to modulate SMD behavior (Figure 5B and Figure 6F). First, sNPF functions only in a neuronal population to elicit SMD behavior. Second, sNPF expression in neurons expressing its receptor is also necessary for SMD behavior. Third, the sNPF expression in PDF-positive neurons that function as the central pacemaker is dispensable for SMD behavior. Fourth, CRY-positive dorsal clock neurons and DN1 clock neurons are essential for sNPF signaling to induce intact SMD behavior. We suggest that the function of CLK/CYC in these neurons might connect the sNPF signaling to modulate timing-based decision making behavior. Fifth, sNPF signaling is necessary both in ellipsoid bodies and mushroom bodies, but not in the fan-shape bodies to elicit memory formation. These results suggest that there is a specialized subset of neurons that expresses sNPF, which specifically contributes to inducing SMD behavior.

We found interesting features by interfering with the function of the sNPF receptor sNPFR1 in precisely defined circuits (Figure 5C and Figure 6G). First, the expression of sNPFR1 both in neuronal and glial population is important to elicit SMD behavior. Second, the expression of sNPFR1 in sNPF-positive cells is dispensable for intact SMD behavior. Third, among clock neurons, PDF-positive central pacemaker cells and DN1 clock cells are dispensable. However, some of the CRY-positive neurons are important for SMD behavior properly. Fourth, sNPF signaling is necessary in all the tested memory circuits such as ellipsoid body, mushroom body and fan-shaped body.

Knocking down sNPFR1 in fan-shaped body neurons diminished SMD behavior. However, inactivation of fan-shaped body neurons using UAS-Kir2.1 had no effect. One scenario compatible with both findings is that sNPFR1 indirectly modulate the function of fan-shaped body neurons. The mechanism of modulating neuronal activity by neuropeptides can be classified in two different time scales. In short-term effects, neuropeptides can modify the activity of ion channels and alter the release of neurotransmitters or the synaptic response. In long-term effects, neuropeptides may signal by altering gene expression and strengthening synaptic transmission via regulating synaptic structures (Chen and Ganetzky, 2012).

It is also worth noting that the sNPFR1-expressing FB cells are not restricted to neuronal cell types. We speculate that glial cell population in this region could be substantial circuit participants to induce SMD behavior through sNPF signaling since sNPFR1 knock down in glial cell population also eliminates this behavior (repo in Figure 5D). However, we cannot exclude the possibility that other as yet unrecognized types of sNPFR1-positive FB cells are involved in SMD behavior as well.

The ability of sNFP knowdown but not Kir2.1 expression in FB cells to impact SMD behavior could also indicate that the sNFP mediated inhibition not activation of fan-shape body neurons is crucial to SMD behavior. If the fan-shape body neurons responsible for SMD behavior are inhibitory neurons, forced depolarization of fan-shape body neurons will disrupt SMD behavior. To test this possibility, we expressed bacterial sodium channel *UAS-NachBac* to force fan-shaped body neurons into a depolarized state. Indeed, such ectopic activation of fan-shaped body neurons disrupted SMD behavior (Figure S4D). In summary, we suggest that sNPF signaling might reduce the neuronal activity of fan-shaped body neurons to induce intact SMD behavior.

As described above, we identified the neural circuits involving sNPF-positive and sNPFR1-positive neurons. In the case of sNPFR1-positive cells, we suggest that some of the glial cell population is required to elicit intact SMD behavior. We also identified the sexually dimorphic FRU-positive sNPF-expressing neurons that modulate this specific behavior (Figure 6). These sexually dimorphic sNPF-expressing neurons show synaptic plasticity depending on the male’s sexual experience (Figure 7A-D and Figure S7B-E). Finally, these neurons alter the neuronal activities following the sexual experiences (Figure 7E-F and Figure S7F-G).

In this paper, we identified the new function of the sNPF in modulating timing-based decision-making process of a male fruit fly and we also found that circadian clock genes Clock/cycle are specifically involved with this process. These results raise the question as to how the sNPF signaling pathway is connected to clock pathways to modulate timing-based decision-making.

### How are Gr5a-positive gustatory inputs connected to sNPF signaling and the CLK/CYC clock networks to modulate mating duration?

We have uncovered contact chemoreception by several male-specific gustatory neurons as a major source for gauging the male fruit fly sexual experience. We found that a male’s sexual fulfillment is provided via gustatory but not olfactory inputs (Figure 2). The fly gustatory system appears to have a simple map compared to the olfactory system, which recognizes and distinguishes thousands of odors, suggesting that the fly may not need to finely distinguish many different tastes. There are two non-overlapping neural populations identified by either Gr5a or Gr66a expression. Gr5a cells recognize sugars and mediate acceptance/attractive behaviors, whereas Gr66a cells recognize bitter compounds and mediate avoidance behavior. In this respect, Gr5a cells can be designated as “acceptance” cells rather than “sweet” cells (Wang, Singhvi, Kong and Scott, 2004). Thus, the specific involvement of Gr5a but not Gr66a cells in SMD is consistent with the notion that both sugar and females will be categorized as acceptance signals. We also found that male-specific Gr5a cells detect the female presence and send signals to the central brain to induce SMD behavior (Figure 3). We then identified LUSH, the odorant binding protein, and SNMP1, the regulator of the pheromone receptor’s signaling in Gr5a-positive cells, as critical components for eliciting the SMD behavior (Figure 2). Thus, we propose that the expression of *snmp1* and *lush* in these Gr5a-positive cells can signal the existence of females via gustatory circuits.

In summary, we found that Gr5a-dependent contact chemoreception is involved in detecting female presence and connected to the sNPF signaling pathway (Figure S5). How are these two circuits connected? First, sNPF signaling has been implicated in regulating feeding behavior in *Drosophila,* similar to its mammalian counterpart NPY (Lee, Kwon, Lee, Kwon, Min, Jung, Kim, You, Tatar and Yu, 2008b). Moreover, indirect evidence suggests that sNPF signaling is required to modulate innate behavioral states such as satiety, which can be shaped by gustatory stimuli to alter neural circuits (Ci, Wu and Su, 2014). Future studies will reveal how Gr5a-positive sensory neurons are connected to sNPF-positive neural circuits. Uncovering the feedback interactions between Gr5a-positive cells and sNPF signaling circuits will shed light on how the information from the sensory modalities can be transmitted to the neuropeptidergic system and integrated into changing internal states of brain dynamics.

There are several lines of evidence indicating that circadian clock genes, especially *Clk* and *cyc*, regulate feeding (Chatterjee, Tanoue, Houl and Hardin, 2010; Krupp and Levine, 2010; Sarov-Blat, So, Liu and Rosbash, 2000; Xu, Zheng and Sehgal, 2008). For example, the input of neuronal clocks and clocks in metabolic tissues are coordinated to mediate effective energy homeostasis with gustatory inputs via *Clk* and *cyc* (Xu, Zheng and Sehgal, 2008). The clock tunes the gustatory system to a higher gain level in the morning that allows the fly to temporally couple the morning bout of activity with food-detection machinery (Chatterjee, Tanoue, Houl and Hardin, 2010). Interestingly, *cyc* function supplied via the *Gr5a-GAL4* driver can rescue the food-detection deficit, which suggests that the perceived meaning of a sensory input is determined not just by the modality or intensity of the stimulus, but also by the circadian time when the stimulus is registered (Chatterjee, Tanoue, Houl and Hardin, 2010). In addition, it was reported that *Clk* and *cyc* specifically regulate starvation-induced sleep-loss, indicating a close relationship between CLK/CYC function and gustatory-dependent metabolic homeostasis (Keene, Duboue, McDonald, Dus, Suh, Waddell and Blau, 2010).

The relationship between sNPF signaling and CLK/CYC function could derive from their common link to the insulin-signaling pathway. Insulin signaling regulates growth, metabolism, and aging of animals, including *Drosophila melanogaster*. *Drosophila* sNPF signaling regulates growth by ERK-mediated insulin signaling (Lee, Kwon, Lee, Kwon, Min, Jung, Kim, You, Tatar and Yu, 2008b). Insulin also interacts with the sNPF signaling pathway by acting as a satiety signal to decrease sNPFR expression, and in turn decrease motivated feeding (Root, Ko, Jafari and Wang, 2011). *Drosophila* FOXO is known to regulate metabolism and oxidative stress. FOXO regulates insulin activity through feedback mechanisms, and also affects circadian rhythms (Zheng, Yang, Yue, Alvarez and Sehgal, 2007). FOXO is expressed predominantly in the fat body, suggesting that *foxo* has non-cell-autonomous effects on central circadian clock function. We propose a possible role of sNPF connecting FOXO-mediated insulin signaling to central and peripheral clock networks. In support of this notion, a previous study demonstrates that the gustatory-dependent metabolic state of a fly influences the clock to modulate behavioral rhythms via insulin and sNPF signaling pathways (Erion and Sehgal, 2013). Finding the possible links among these circuits would be an interesting subject for future investigation.

### Neural reuse model: how the same clock neurons modulate both circadian and interval timing-based behaviors

It is common for neural circuits established for one purpose to be exapted during evolution (exaptation) (Gould and Vrba, 1982; Skoyles, 1999) for different uses without losing their original function. ‘Neural reuse’ theories can explain the evolutionary-developmental properties of neural plasticity involved in human cognition such as reuse of motor control circuits for language, reuse of motor control circuits for memory, and reuse of circuits for numerical cognition (Anderson, 2010). Neural reuse may be more prevalent in insect brains since many insects possess small brains that have been miniaturized during evolution. Their small size suggests that insects are under selective pressure to reduce energetic costs and brain size. Thus, in smaller brains, there may be an increased prevalence of neural reuse (Chittka and Niven, 2009; Niven and Chittka, 2010).

In support of this hypothesis, it has been proposed that dCLK/CYC-controlled transcriptional/translational feedback loops operate differently in subsets of pacemaker neurons (Lee, Cho, Kang, Jeong, Chen, Yoo and Kim, 2016). Thus the same CLK/CYC-controlled molecular pathways may contribute to the specific functions dependent on the different neural circuits. Consistent with this hypothesis, we previously showed that male flies use a subset of PDF/NPF-expressing clock neurons, which were originally established to regulate circadian rhythm, for processing the information of previous exposure to rivals (Kim, Jan and Jan, 2013). Here we propose another evidence of neural reuse model found in the fly brain to modulate both circadian and interval timing-based behaviors. Our studies suggest that neural reuse occurs in the brain of a fruit fly similar to that of humans (Anderson, 2010; Dumont and Robertson, 1986; Skoyles, 1999).

### Possible relationship between SMD and LMD

Our earlier study found that previous exposure to rivals lengthens mating duration, a behavior named Longer-Mating-Duration (LMD) (Kim, Jan and Jan, 2012). Even though LMD and SMD share a common behavioral phenotype in regulating mating duration, there are clear differences in the underlying mechanisms (Table 1—Move this table from supplemental material to the main text as it nicely compare LMD with SMD). First, LMD requires only visual input while SMD requires multiple inputs, including gustatory input. Second, LMD requires the function of the clock genes PER/TIM, whereas SMD requires the function of CLK/CYC. Third, LMD requires the EB for its memory storage, but SMD requires both the EB and MB. Fourth, *rut* and *amn* are important to process the memory for LMD, while *orb2* is critical for SMD. Fifth, LMD requires the function of sexually dimorphic LNd clock neurons, however SMD requires that of Gr5a-positive sensory neurons. Thus both behaviors are involved in modulating the length of copulation but use different genetic components and different but partially overlapping neuronal circuitry.

These two behavioral circuits might have evolved independently since they use different sensory cues for detecting ‘rivals vs. females’ for ‘sexual competition vs. sexual satiety’. Male flies depend on visual input to recognize rivals (Kim, Jan and Jan, 2012), however they depend on female pheromones via contact chemoreception to recognize female presence and register sexual experience (Figures 1 and 2). Recent studies demonstrating the need of males to touch the female body in order to persistently induce female-oriented behaviors also support this premise (Agrawal, Safarik and Dickinson, 2014; Kohatsu and Yamamoto, 2015). We propose that contact chemoreception induced by female body pheromones contributes to the determination of a male’s mating investment in *Drosophila melanogaster*. Like LMD (Kim, Jan and Jan, 2012), SMD requires clock genes involved in non-circadian functions, such as learning and memory, habituation, sleep, drug sensitization, and mating duration (Hall, 2005), without the feedback between PER/TIM and CLK/CYC that is crucial for circadian rhythm.

Even though LMD and SMD use different sensory modalities and neuropeptide expressing neurons, that information needs to be collected together and sent to output neurons for the termination of mating. It has been reported that mating termination involves neuropeptide corazonin (Crz) expressing neurons, subset of dopaminergic and GABAergic neurons (Crickmore and Vosshall, 2013; Tayler, Pacheco, Hergarden, Murthy and Anderson, 2012). Further investigation will be required to find possible connections between LMD and SMD circuits with the output neural circuitry.

### Both SMD and LMD provide a social behavior model using fruit fly

As noted by Darwin, the brain of social insects is the most marvelous atoms of matter in the world more so than the brain of man (Darwin, 1871), considering how the remarkably complex neural circuits which process social information are integrated into such a tiny space (Chittka and Niven, 2009). Unlike the classic eusocial insects such as ants and bees, flies are not typically regarded for their group dynamics, however, this view is changing with growing evidence of social interactions in *Drosophila* (Battesti, Moreno, Joly and Mery, 2012; Benzer, 1967; Billeter, Jagadeesh, Stepek, Azanchi and Levine, 2012; Billeter, Atallah, Krupp, Millar and Levine, 2009; Billeter and Levine, 2013; Bretman, Fricke and Chapman, 2009; Ganguly-Fitzgerald, Donlea and Shaw, 2006; Garbaczewska, Billeter and Levine, 2013; Kacsoh, Bozler, Hodge, Ramaswami and Bosco, 2015; Kacsoh, Bozler, Ramaswami and Bosco, 2015; Kent, Azanchi, Smith, Formosa and Levine, 2008; Kim, Jan and Jan, 2012, 2013; Krupp, Kent, Billeter, Azanchi, So, Schonfeld, Smith, Lucas and Levine, 2008; Krupp and Levine, 2010; Lefranc, Jeune, Thomas-Orillard and Danchin, 2001; Levine, Funes, Dowse and Hall, 2002; Lewis, Heys, Prescott and Lizé, 2014; Loyau, Blanchet, Van Laere, Clobert and Danchin, 2012; Mery, Varela, Danchin, Blanchet, Parejo, Coolen and Wagner, 2009; Saltz and Foley, 2011; Sarin and Dukas, 2009; Schneider, Dickinson and Levine, 2012; Tinette, Zhang and Robichon, 2004; Vosshall, 2007; Wang and Anderson, 2010; Wang, Han, Mehren, Hiroi, Billeter, Miyamoto, Amrein, Levine and Anderson, 2011a). Moreover, a recent study suggests that *Drosophila melanogaster* exhibits collective behavior that has been investigated in schooling fish, flocking birds and human crowds (Ramdya, Lichocki, Cruchet, Frisch, Tse, Floreano and Benton, 2015). It is clear that social communication is actually much more fundamental to life than previously considered (Kacsoh, Bozler, Ramaswami and Bosco, 2015). Thus, the fly is a useful model system as a compromise between tractability and richness, to study the conserved neural circuits that elicit social behavior (Burne, Scott, van Swinderen, Hilliard, Reinhard, Claudianos, Eyles and McGrath, 2011; Olsen and Wilson, 2008; Vosshall, 2007). Together with recent studies of social experience-mediated and context-dependent behavior of fruit flies, LMD and SMD will help us understand how animals receive social information and then process it at the molecular level.

### SMD behavior provides a novel and genetically traceable satiety paradigm

Satiety is usually defined as the state of being fed beyond capacity. Along this line, satiety studies were focusing on feeding behaviors in *Drosophila* (Krashes, DasGupta, Vreede, White, Armstrong and Waddell, 2009; Nassel and Williams, 2014; Schleyer, Saumweber, Nahrendorf, Fischer, von Alpen, Pauls, Thum and Gerber, 2011; Söderberg, Carlsson and Nässel, 2012). In this context, the distinctive feature of satiation defined as the blunted sensitivity to food tastes, especially to sugars. In flies, myoinhibitory peptide (MIP) has been identified as an important factor to coordinate feeding motivation and maintain a constant body weight (Min, Chae, Jang, Choi, Lee, Jeong, Jones, Moon, Kim and Chung, 2016; Min and Chung, 2016). The drosulfakinin (DSK) peptide has been identified as satiety signals together with insulin-like peptides (DILPs) (Soderberg, Carlsson and Nassel, 2012). DSK is a cholecystokinin-like peptide found in the fruit fly, and cholecystokinin signaling appears well conserved during evolution. Like sNPF, DSK is a multifunctional neuropeptide that regulates gut function, satiety and food ingestion, hyperactivity and aggression, and synaptic plasticity during neuromuscular junction and development (Nassel and Williams, 2014).

Sexual satiety defines the long-term inhibition of masculine sexual behavior after repeated ejaculations in rats and mice. By measuring the c-Fos expression level in the brain regions of the sexually satiated male, researchers identified several areas in the male rat forebrain of relevance to sexual satiety (Phillips-Farfan and Fernandez-Guasti, 2007). Recently Zhang and colleagues reported that *Drosophila* males show reproductive satiety (Zhang, Rogulja and Crickmore, 2016). In their behavioral paradigm, males’ mating drive gets reduced after 4 hours of exposure to excessive females. The major difference between this report and ours is that Zhang and colleagues did not specifically measure the mating duration. Instead, they measured all the mating behaviors during the 4 hours of time frame and interpreted the reduced mating performance as satiety index. DopR1-mediated superior medial protocerebrum (SMPA) dopaminergic circuits has been identified as an important element for the mating drive.

Here we demonstrate novel forms of satiety paradigm related to sexual experience, not to food ingestion. We found that sexually satiated male *Drosophila* reduces the investment on mating (Figure 1). Surprisingly, gustatory neurons crucial for sugar detection also detect female pheromone to register the satiety states in the central brain (Figure 2). Since ILP, DSK, and MIP have been reported as modulators of food-related satiety signaling, future study will investigate the relationship between the brain circuitry we found and the ILP/DSK circuits (Min, Chae, Jang, Choi, Lee, Jeong, Jones, Moon, Kim and Chung, 2016; Min and Chung, 2016; Nassel and Williams, 2014; Soderberg, Carlsson and Nassel, 2012). This investigation will help us identify the fundamental principles regarding how food ingestion-related satiety and sexual satiety signaling share the neural circuits and still maintain their respective independent functionality. It will also be interesting to investigate the role of dopaminergic circuits in SMD behavior since dopamine plays a crucial role to modulate reproductive satiety of male fruit fly (Zhang, Rogulja and Crickmore, 2016).

In summary, we report a novel anorexigenic pathway that controls satiety related to sexual experiences in *Drosophila*. This novel paradigm will provide a new avenue to study how the brain actively maintains a constant mating drive to invest on sexual behavior properly.

## ACKNOWLEDGEMENTS

We thank Dr. Kweon Yu (Korea Research Institute of Bioscience and Biotechnology: KRIBB, Korea) for kindly providing valuable sNPF-related fly strains. We thank Dr. Joshua Bagley (UCSF) and Dr. Kyeongjin Kang (Sungkyungkwan University, Korea) for helpful comments on this paper. The work was supported by NIH grant 2R37NS040929 to YNJ. LYJ and YNJ are investigators of Howard Hughes Medical Institute.

## EXPERIMENTAL PROCEDURES

### Fly Rearing and Strains

*Drosophila melanogaster* were raised on cornmeal-yeast medium at similar densities to yield adults with similar body sizes. Flies were kept in 12 h light: 12 h dark cycles (LD) at 25°C (ZT 0 is the beginning of the light phase, ZT12 beginning of the dark phase) except for some experimental manipulation (experiments with the flies carrying *tub-GAL80^ts^*). Wild-type flies were *Canton-S*. To reduce the variation from genetic background, all flies were backcrossed for at least 3 generations to CS strain. All mutants and transgenic lines used here have been described previously.

We are very grateful to the colleagues who provided us with many of the lines used in this study. We obtained the following lines from Dr. Joel D. Levine and Joshua J. Krupp (University of Toronto, Canada): *PromE(800)-GAL4* (*oeno-GAL4* in this study); from Dr. Kweon Yu (Korea Research Institute of Bioscience and Biotechnology: KRIBB, Korea), *sNPF-GAL4*, *UAS-sNPF*, *sNPF-RNAi*, *sNPF-R-RNAi*; from Dr. Barry Dickson (HHMI Janelia Research Campus, USA): *UAS[stop]mCD8GFP; fru^FLP^, UAS[stop]DscamGFP; fru^FLP^, UAS[stop]nsybGFP; fru^FLP^, UAS[stop]TNT_active_; fru^FLP^, UAS[stop]TNT_inactive_; fru^FLP^, UAS[stop]RicinA; fru^FLP^*, *fru-GAL4, orb2^Δ^, orb2^ΔQ^*; from Dr. Chung-Hui Yang (Duke University Medical Center, USA): *Gr5a-GAL4*, *elav-GAL80*; from Dr. Toshiro Aigaki (Tokyo Metropoitan University, Japan): *UAS-mSP*; from Dr. Herman Wijnen (University of Virginia, USA) (Goda, Mirowska, Currie, Kim, Rao, Bonilla and Wijnen, 2011): *UAS-cyc; cyc^01^, elav^c155^; UAS-cyc; cyc^01^, elav^c155^; pdf-GAL80; cyc^01^, elav^c155^; tub-GAL80^ts^; cyc^01^;* from Dr. Martin Heisenberg (Universität Würzburg, Germany): *WT Berlin*, *ninaE^17^*; from Dr. Alex C. Keene (University of Nevada, USA) and Dr. Justin Blau (New York University, USA): *cyc^01^, Clk^Jrk^, cry-GAL80;cry-GAL80, tim-GAL4, pdf-GAL4*, *cry-GAL4*; from Dr. Ravi Allada (Northwestern University, USA): *Clk4.1M-GAL4*; from Dr. Amita Sehgal (University of Pennsylvania Medical School, USA: *pdf^01^*, *per^01^, tim^01^, per^01^;tim^01^*; from Dr. J. Douglas Armstrong (University of Edinburgh): *c547-GAL4*; from Dr. Jing Wang (University of California San Diego, USA): *UAS-mLexA-VP16-NFAT, LexAop-CD2-GFP; LexAop-CD8-GFP-2A-CD8-GFP*; from Dr. Gero Miesenböck (University of Oxford, UK): *MB-GAL80*; from Dr. Ulrike Heberlein (HHMI Janelia Research Campus): *14-94-GAL4*; Dr. Chun Han and Dr. Peter Soba generated the following lines from our laboratory: *UAS-CD4-tdGFP, UAS-CD4-tdTomato.*

The following lines were obtained from Bloomington Stock Center (#stock number): *Orco^1^* (#23129)*, Orco^2^* (#23130*), dnc^1^* (#6020*), rut^2080^* (#9405*), amn^1^* (#5954), *UAS-tubGAL80^ts^* (#7018*), UAS-Kir2.1* (#6596), *UAS-NachBac* (#9469), *Df(1)Exel6234* (#7708)*, UAS-tra^F^* (#4590), *GustD^x6^* (#8607), *Gr66a-GAL4* (#28801), *UAS-mCD8GFP* (#5130), *UAS-RedStinger* (#8547), *snmp1^1^* (#25043), *snmp1^2^* (#25042), *UAS-snmp1* (#25044), *UAS-TNT* (#28997), *sNPF-R-GAL4* (#46547), *npfR1^c01896^* (#10747), *Df(3R)BSC464* (*npfR1* deficiency line, #24968), *sNPF-R^MI00427^* (#30996), *elav^c155^* (#458), *repo-GAL4* (#7415), *UAS-dicer* (#24650, #24651), *UAS-Denmark; UAS-syteGFP* (#33064), *UAS-n-sybGFP* (#6921), *npf-GAL4* (#25682), *tra-RNAi* (#28512), *lush-RNAi* (#31657), *orb2-RNAi* (#27050); from Kyoto Stock Center (DGRC): *iav^1^* (#101174), *ok107-GAL4* (#106098), *GMR-Hid* (#108419); from Vienna Stock Center (VDRC*): Clk-RNAi* (#v42833, #v42834), *cyc-RNAi* (#v11765); from Harvard Exelixis Collection: *sNPF^c00448^* (#c00448).

### Mating Duration Assays

Mating duration assay was performed as previously described (Kim, Jan and Jan, 2012). For naïve males, 4 males from the same strain were placed into a vial with food for 5 days. For experienced males, 4 males from the same strain were placed into a vial with food for 4 days then eight CS virgin females were introduced into vials for last 1 day before assay. Five CS females were collected from bottles and placed into a vial for 5 days. These females provide both sexually experienced partners and mating partners for mating duration assays. At the fifth day after eclosion, males of the appropriate strain and CS virgin females were mildly anaesthetized by CO_2_. After placing a single female in to the mating chamber, we inserted a transparent film then placed a single male to the other side of the film in each chamber. After allowing for 1 h of recovery in the mating chamber in a 25°C incubator, we removed the transparent film and recorded the mating activities. Only those males that succeeded to mate within 1 h were included for analyses. Initiation and completion of copulation were recorded with an accuracy of 10 sec, and total mating duration was calculated for each couple. All assays were performed from noon to 4 pm.

### Courtship assays

Courtship assay was performed as previously described (Ejima and Griffith, 2007), under normal light conditions in circular courtship arenas 11 mm in diameter, from noon to 4 pm. Courtship latency is the time between female introduction and the first obvious male courtship behavior such as orientation coupled with wing extensions. Once courtship began, courtship index was calculated as the fraction of time a male spent in any courtship-related activity during a 10 min period or until mating occurred. Mating initiation is the time after male flies successfully mounted on female.

### Locomotion assays

For climbing assay, individual flies were placed in a 15 ml falcon tube (Fisher Scientific) and were gently tapped to the bottom of the tube. The time taken for the flies to climb 8 cm of the tube wall was recorded. Each fly was tested 5 times. Other than a single instance, all flies were seen to reach the target height within 2 min, which was the experimental cut-off time. Velocity was obtained by dividing the lines (mm) a fly crossed (distance walked) by time (sec) a fly reached the line of the tube. For horizontal (spontaneous) locomotor activities, a single fly was first brought to the middle of the column by gentle shaking and then the fly movement was constantly monitored for 5 min and recorded. Total fraction of time flies walked during 5 min was calculated and number of stops during 5 min was also counted then calculated (Mohammad, Singh and Sharma, 2009).

### Immunostaining and antibodies

As described before (Lee and Luo, 1999), brains dissected from adults 5 days after eclosion were fixed in 4% formaldehyde for 30 min at room temperature, washed with 1% PBT three times (30 min each) and blocked in 5% normal donkey serum for 30 min. The brains were then incubated with primary antibodies in 1% PBT at 4°C overnight followed with fluorophore-conjugated secondary antibodies for 1 hour at room temperature. Brains were mounted with anti-fade mounting solution (Invitrogen, catalog #S2828) on slides for imaging. Primary antibodies: chicken anti-GFP (Aves Labs, 1:1000), rabbit anti-DsRed express (Clontech, 1:250), mouse anti-Bruchpilot (nc82) (DSHB, 1:50), mouse anti-PDF (DSHB, 1:100). Fluorophore-conjugated secondary antibodies: Alexa Fluor 488-conjugated goat anti-chicken (Invitrogen, 1:100), Alexa Fluor 488-conjugated donkey anti-rabbit (Invitrogen, 1:100), RRX-conjugated donkey anti-rabbit (Jackson Lab, 1:100), RRX-conjugated donkey anti-mouse (Jackson Lab, 1:100), Dylight 649-conjugated donkey anti-mouse (Jackson Lab, 1:100).

### Imaging and quantitative analysis of GFP Fluorescence

As described before (Kayser, Yue and Sehgal, 2014a), brains were visualized with a TCS SP2 confocal microscope and images processed with ImageJ (National Institutes of Health, Bethesda, MD; <u>http://rsb.info.nih.gov/ij</u>). All settings were kept constant between experimental conditions. Images were taken in 0.5 μM steps unless otherwise specified. For CaLexA experiments, GFP fluorescence for CaLexA was normalized to nc82 staining, and then the fluorescence of ROI was quantified using the histogram tool of ImageJ. Both hemispheres of six fly brains were analyzed (total 12) for statistical analysis. For quantifying *n-sybGFP* particles, we used the “analyze particles” function of ImageJ. Average size represents the average size of analyzed particles. Percent of area represents percent of area that is covered with particles normalized by total area. All imaging and analysis were done blind to experimental condition.

### Statistical Analysis

Statistical analysis of mating duration assay was described previously (Kim, Jan and Jan, 2012). More than 36 males (naïve or experienced) were used for mating duration assay. Our experience suggests that the relative mating duration differences between naïve and experienced condition are always consistent; however, both absolute values and the magnitude of the difference in each strain can vary. So we always include internal controls for each treatment as suggested by previous studies (Bretman, Westmancoat, Gage and Chapman, 2011). Therefore, statistical comparisons were made between groups that were naively reared or sexually experienced within each experiment. As mating duration of males showed normal distribution (Kolmogorov-Smirnov tests, p > 0.05), we used two-sided Student’s *t* tests. Each figure shows the mean ± standard error (s.e.m) (*** = p < 0.001, ** = p < 0.01, * = p < 0.05). All analysis was done in GraphPad (Prism). Individual tests and significance are detailed in figure legends.

When we compare the difference of mating duration in experiments without internal control built in (Figure 1C and 1D), we always performed control experiments of wild type for each independent experiment for internal comparison. And in this case, we analyzed data using ANOVA for statistically significant differences (at a 95.0% confidence interval) between the means of mating duration for all conditions. If a significant difference between the means was found by Kruskal-Wallis test, then the Dunn’s Multiple Comparison Test was used to compare the mean mating duration of each condition to determine which conditions were significantly different from condition of interest. (^#^ = p < 0.05)

---

**Supple.Fig.1.**
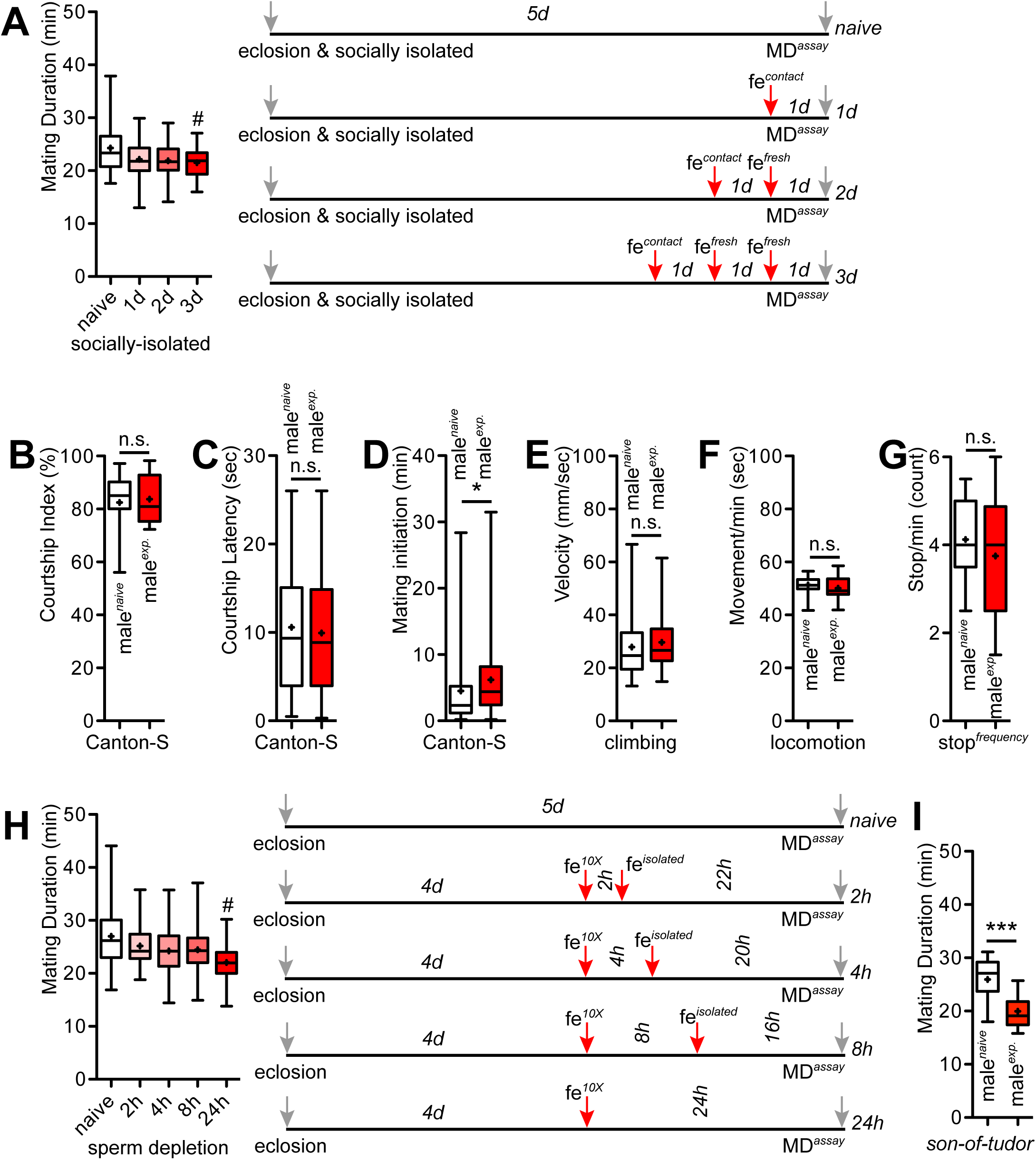

**Supple.Fig.2.**
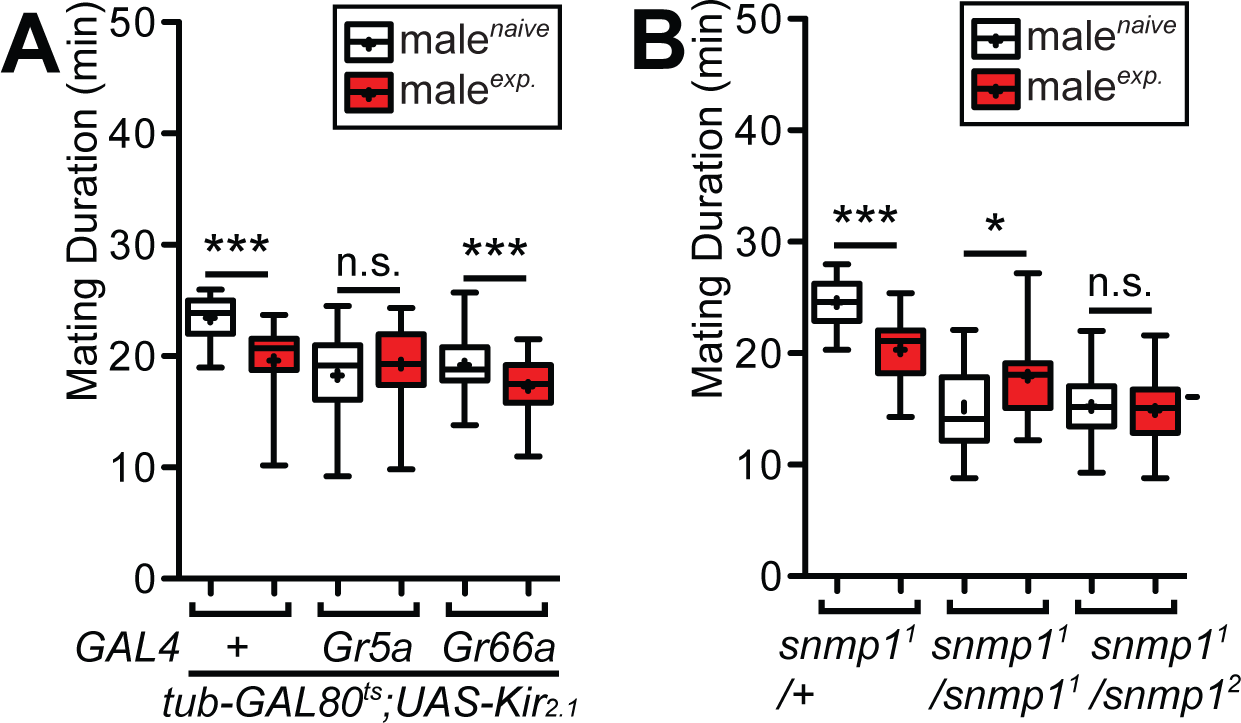

**Supple.Fig.3.**
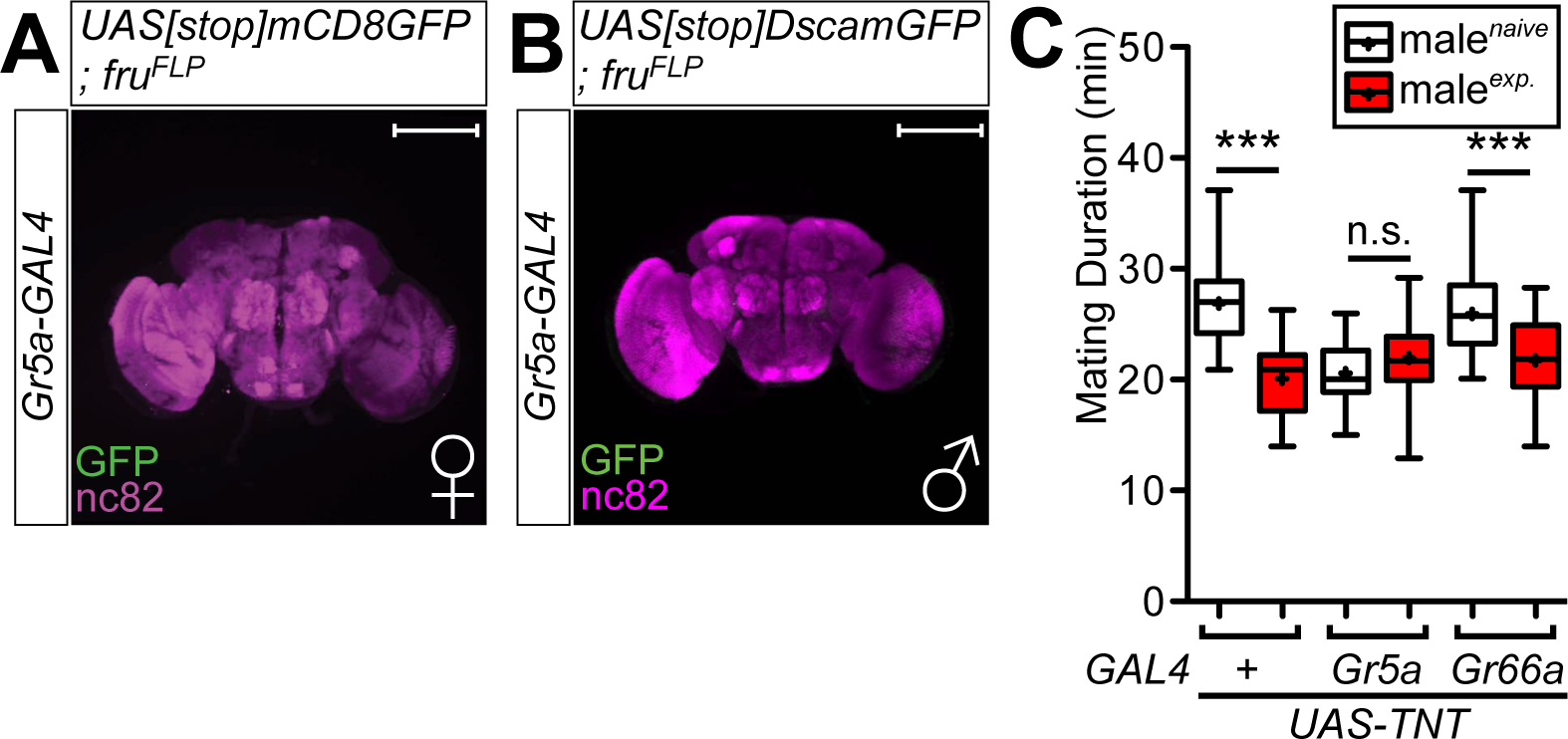

**Supple.Fig.4.**
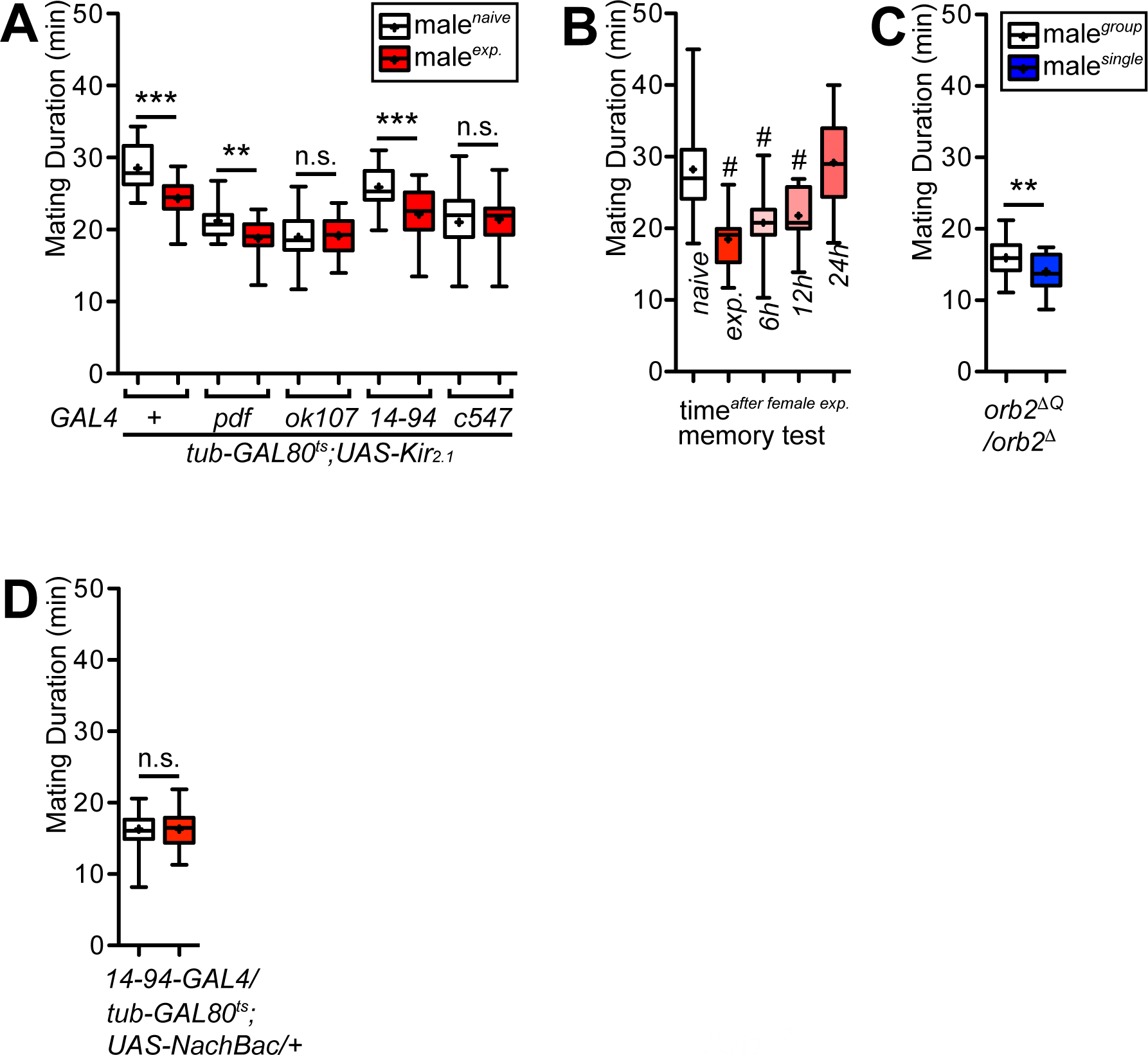

**Supple.Fig.5.**
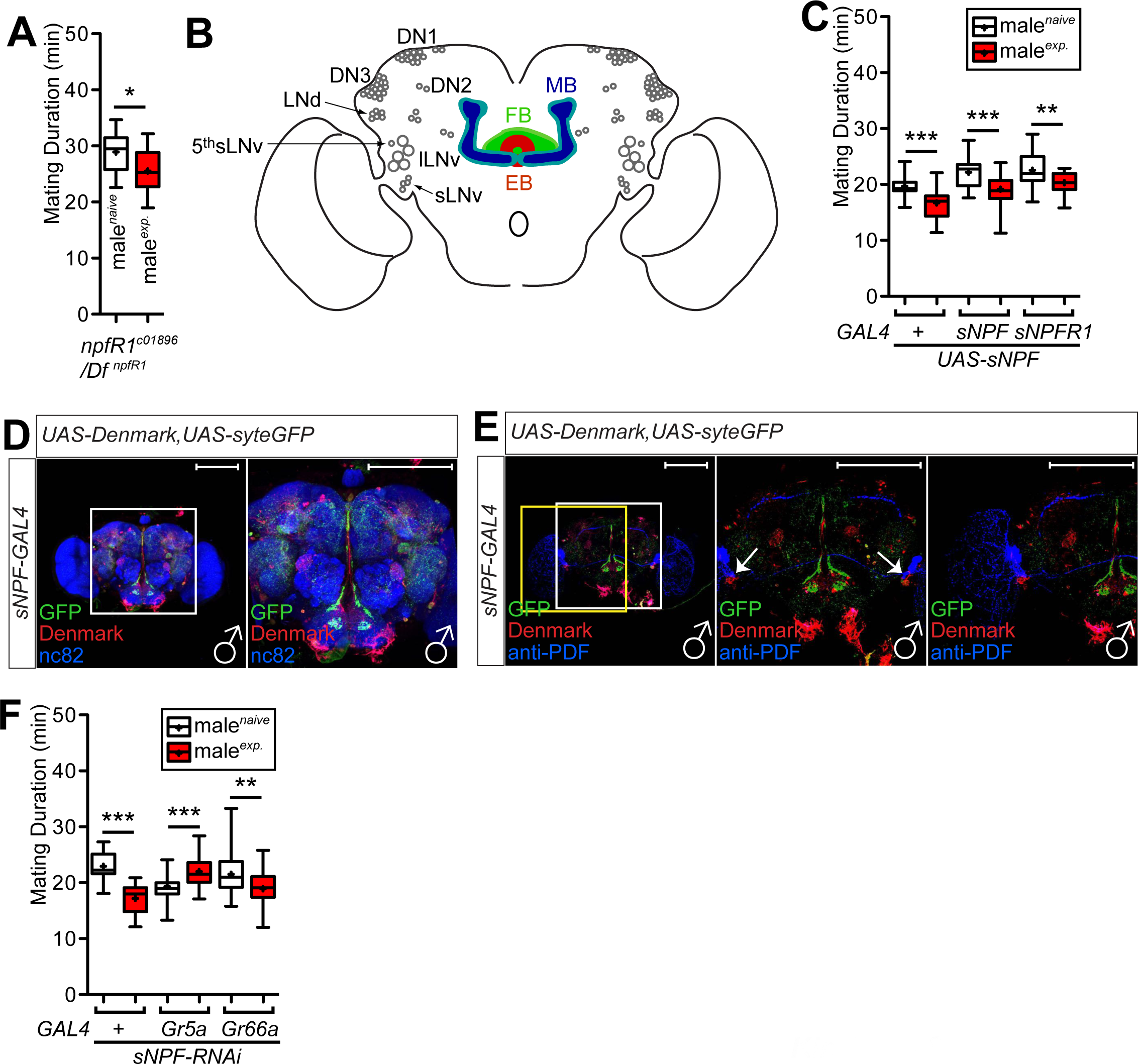

**Supple.Fig.6.**
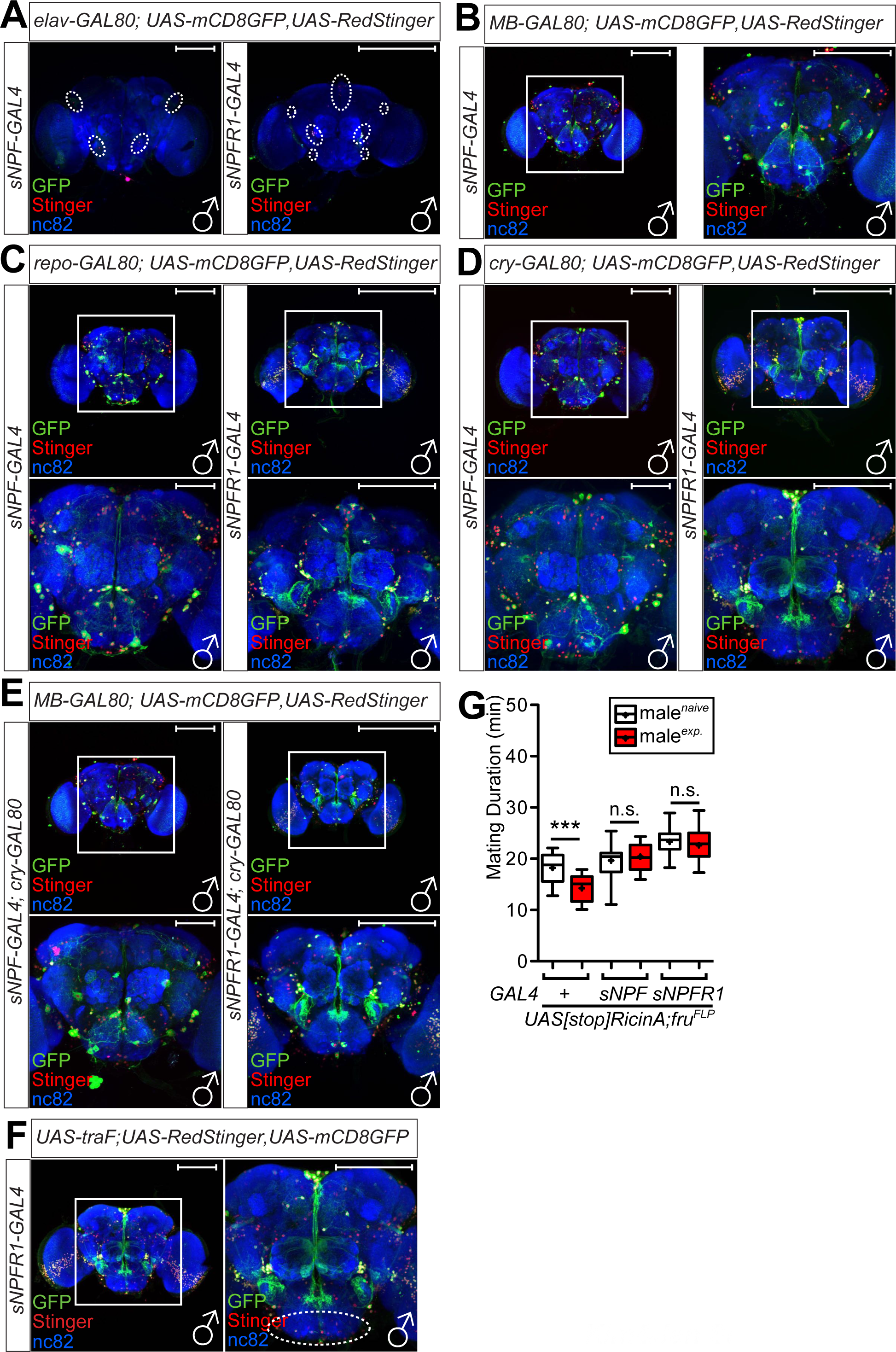

**Supple.Fig.7.**
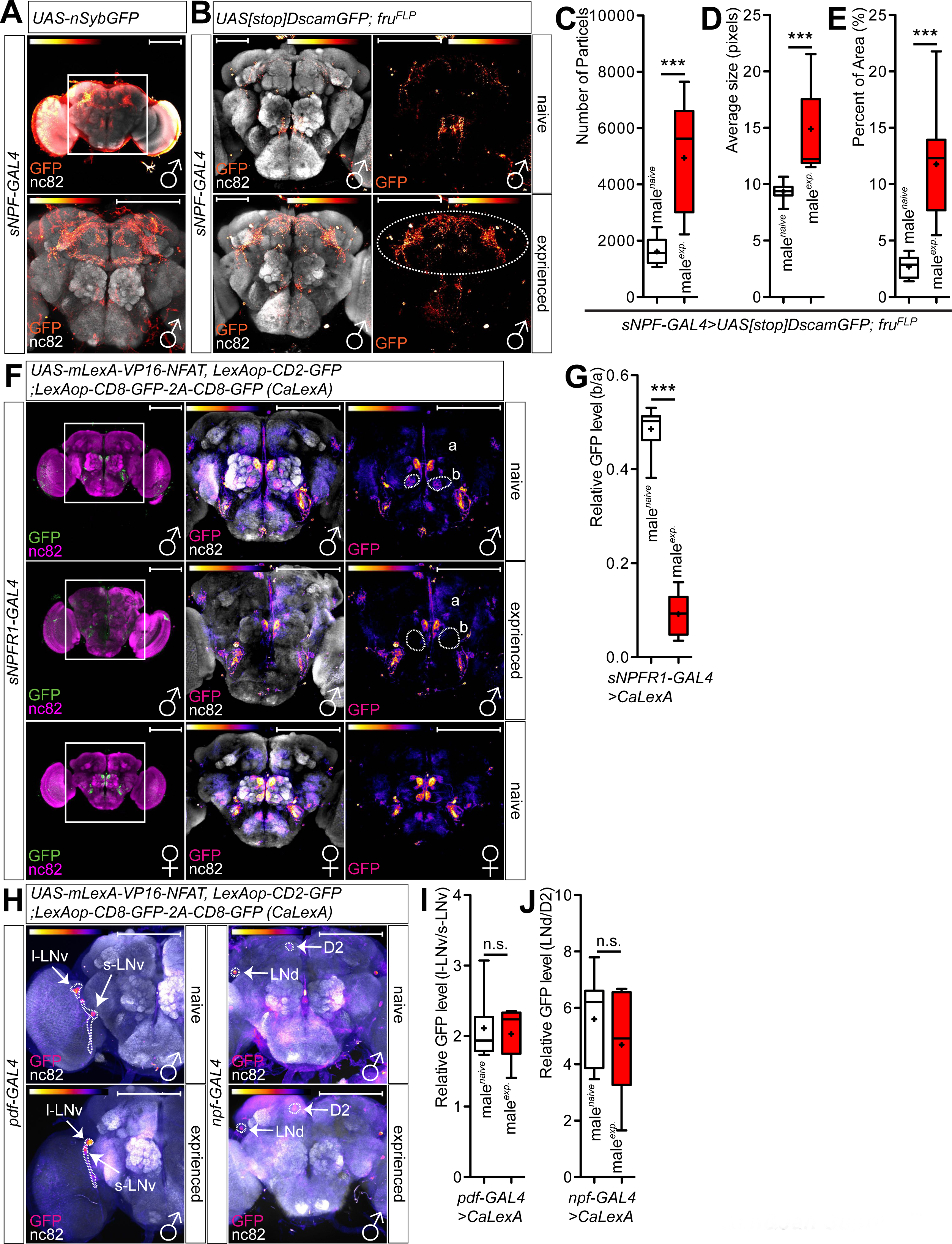

**Supple.Table.1.**

|  | LMD<sup>a</sup> | SMD |
| --- | --- | --- |
| Sensory input | vision | taste, touch |
| Clock components | PER/TIM | CLK/CYC |
| Neuropeptide | PDF/NPF | sNPF |
| Memory circuit (*gene*) | EB (*rut*, *amn*) | EB & MB (*orb2*) |
| Sexual dimorphic cells | 2 LNd clock neurons | ~4 Gr5a taste neurons |

